# The Aspartyl Protease Ddi1 Is Essential for Erythrocyte Invasion by the Malaria Parasite

**DOI:** 10.1101/2021.05.11.443575

**Authors:** Sophie Ridewood, A. Barbara Dirac-Svejstrup, Stephen Howell, Anne Weston, Christine Lehmann, Asha Parbhu Patel, Lucy Collinson, Ryan Bingham, David Powell, Ambrosius Snijder, Jesper Q. Svejstrup, Edgar Deu

## Abstract

Malaria pathology is caused by the exponential replication of *Plasmodium* parasites in the blood stream. The bottleneck of the parasite life cycle is the invasion of erythrocytes immediately after parasites egress from infected red blood cells. DNA damage-inducible protein 1 (Ddi1) is a conserved eukaryotic proteasome shuttle protein containing an internal retroviral-like protease domain. Using conditional genetics, we now show that the proteolytic activity of the *P. falciparum* homologue, PfDdi1, is critically required for invasion of red blood cells. Furthermore, PfDdi1 disruption results in the accumulation of highly polyubiquitinated proteins that can be processed by purified PfDdi1 or distant eukaryotic homologues. We also show that PfDdi1 interacts with multiple components of the ubiquitin-proteasome system and that parasites lacking PfDdi1 are more sensitive to proteasome inhibition. Overall, this study establishes PfDdi1 as a key component of the eukaryotic ubiquitin-proteasome system and as a promising antimalarial target.

## Introduction

Malaria remains one of the most devastating infectious diseases worldwide with more than 200 million clinical cases per year and close to half a million deaths (WHO, 2018). Over the last 15 years, the implementation of global vector control programs and artemisinin-combination therapies have led to a very significant reduction in malaria incidence and mortality (Bhatt et al., 2015). However, the rise of mosquito resistance to insecticides and the emergence and spread of resistance to all frontline antimalarial drugs, including artemisinin-combination therapies, attest to the urgency of identifying novel drug targets and developing new malaria treatments (Tse et al., 2019).

Malaria is caused by *Plasmodium* apicomplexan parasites that are transmitted by female *Anopheles* mosquitoes during a blood meal. Parasites are injected into the epidermis of the human host and travel to the liver where they establish an asymptomatic infection. After replication in hepatocytes, thousands of merozoites are released into the blood stream to initiate the erythrocytic cycle. This consists of invasion of red blood cells (RBCs) by merozoites, forming what is known as ring stage parasites. As these forms grow within the infected RBC (iRBC) they transition into trophozoite stage parasites that occupy most of the RBC volume. This is followed by asynchronous nuclear division (schizont stage) and a final cytokinesis step to form 20-32 daughter merozoites that then egress from the iRBC to start the cycle anew. A small proportion of parasites will develop into male and female gametocytes and will mature into gametes in the midgut of the mosquito after a blood meal. There they will reproduce sexually, replicate and travel to the salivary gland from where they will be transmitted to the next human host. The synchronous exponential growth of parasites during the erythrocytic cycle is responsible for malaria pathology, and therefore, antimalarial drugs need to block parasite replication at this stage. However, in order to have prophylactic effects and block transmission, an ideal drug should also target the liver and sexual stages of parasite development (Burrows et al., 2013).

Proteases play essential roles at all stages of parasite development and are therefore potential antimalarial targets (Deu, 2017). For instance, *Plasmodium* pepsin-like aspartyl proteases are involved in a variety of essential biological processes such as hemoglobin degradation, protein export, RBC invasion or parasite egress. However, there is another putative aspartyl protease that is yet to be characterized in *Plasmodium*, DNA-damage inducible protein 1 (Ddi1). Ddi1 contains an unusual retroviral protease-like (RVP) domain that bears more structural similarity to HIV protease than pepsin-like proteases (Sirkis et al., 2006). In contrast to monomeric pepsin-like aspartyl proteases, RVPs are homodimeric with each monomer providing one of the two catalytic aspartates required for peptide bond hydrolysis.

Ddi1 proteins are conserved across eukaryotes, with all thought to possess an active RVP domain. These proteins have diverse roles in eukaryotes, such as regulation of protein secretion (Lustgarten & Gerst, 1999; Marash & Gerst, 2003; White et al., 2011), response to cellular stress (Kottemann et al., 2018; Serbyn et al., 2019; Svoboda et al., 2019) and activation or degradation of proteins (Ivantsiv et al., 2006; Kaplun et al., 2005; Koizumi et al., 2016; Lehrbach & Ruvkun, 2016). Many of these roles involve interactions with ubiquitin-proteasome system (UPS). One suggested function of Ddi1 is as a proteasomal shuttle factor that binds ubiquitinated cargo and delivers them for proteasomal degradation (Saeki, 2017; Zientara-Rytter & Subramani, 2019). This role is thought to be mediated by additional N-terminal ubiquitin-like (UBL) and C-terminal ubiquitin-associated (UBA) domains or ubiquitin-interacting motifs (UIMs) that allow Ddi1 proteins to bind to proteasomal subunits and ubiquitin (Figure 1A) (Bertolaet et al., 2001; Gomez et al., 2011; Kang et al., 2006; Nowicka et al., 2015; Saeki et al., 2002; Sivá et al., 2016). An increasingly well-characterized role in metazoans is the Ddi1-dependent cleavage and activation of the transcription factor NRF1, which upregulates the expression of proteasomal subunits (Koizumi et al., 2016; Lehrbach & Ruvkun, 2016). As such, there is interest in targeting human DDI2 (hDdi2) alongside the proteasome in cancer therapy in order to prevent the proteasome ‘bounce-back’ response. Indeed, recent studies have shown that knockout (Dirac-Svejstrup et al., 2020) or knock down (Northrop et al., 2020) of hDdi2 renders cancer cells more sensitive to proteasome inhibition, and that HIV protease inhibitors that target hDdi2 have synergistic effects with proteasome inhibitors (Fassmannová et al., 2020; Gu et al., 2020). Progressing these proteins as drug targets has been complicated by a general inability to detect aspartyl protease activity of Ddi proteins *in vitro* (Koizumi et al., 2016; Nowicka et al., 2015; Sivá et al., 2016; Trempe et al., 2016) despite extensive evidence of the requirement for a catalytically active RVP *in vivo* (Gabriely et al., 2008; Koizumi et al., 2016; Lehrbach & Ruvkun, 2016; Svoboda et al., 2019; White et al., 2011). However, recent work has demonstrated that both *Saccharomyces cerevisiae* Ddi1 (ScDdi1) (Yip et al., 2020) and hDdi2 (Dirac-Svejstrup et al., 2020) possess a unique ubiquitin-directed endopeptidase activity, and loss of hDdi2 activity leads to an accumulation of hyper-ubiquitinated proteins, including NRF1 (Dirac-Svejstrup et al., 2020).

**Figure 1.**
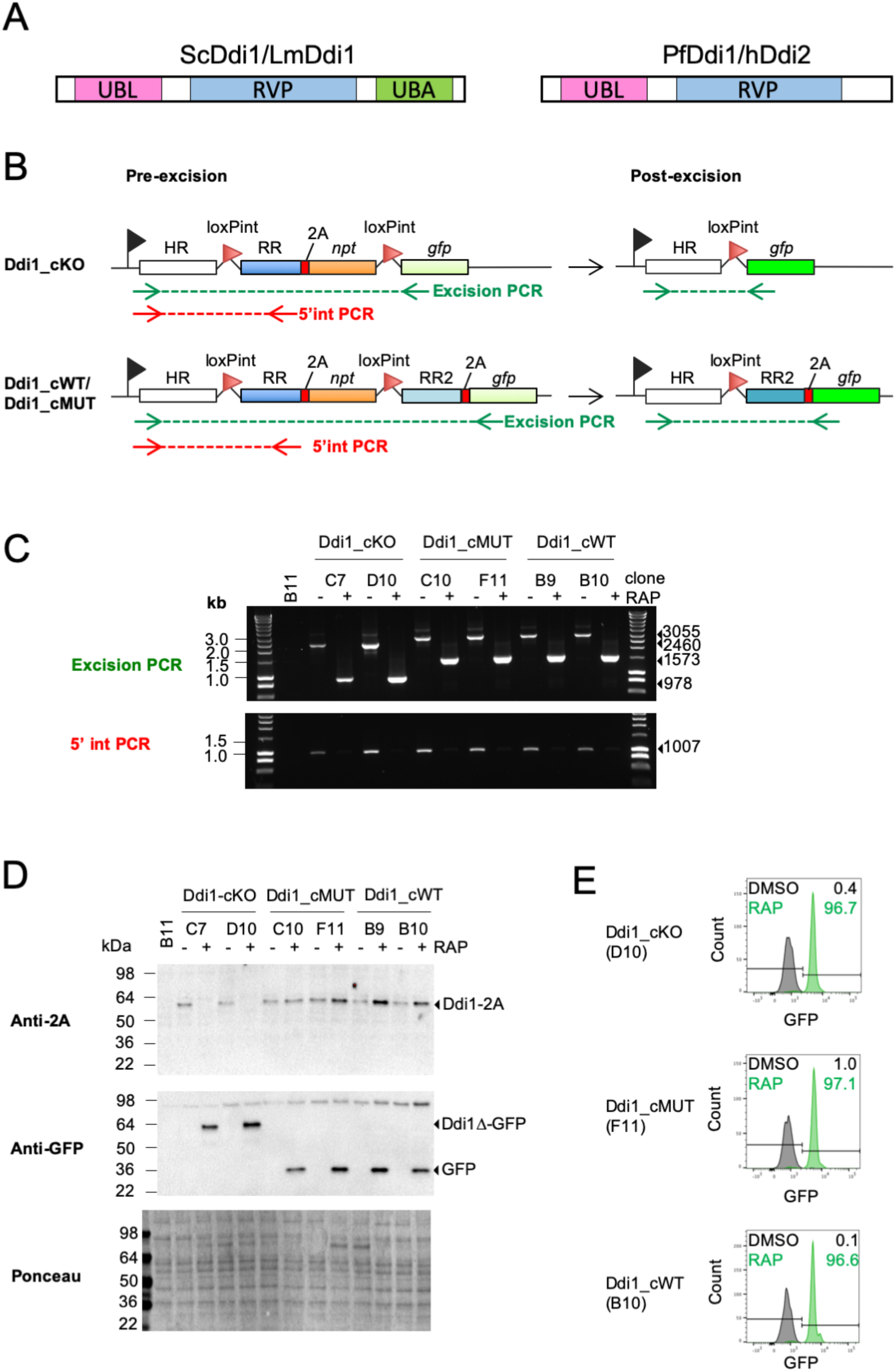
Generation of PfDdi1 conditional lines. (**A**) Domain organization of Ddi1 proteins from different organisms: yeast (ScDdi1), human (hDdi2), *Leishmania* (LmDdi1), and *P. falciparum* (PfDdi1). (**B**) Schematic of the modified *PfDdi1* locus before (left) and after (right) RAP treatment to induce conditional truncation (PfDdi1_cKO, top) and conditional allelic replacement (PfDdi1_cWT & PfDdi1_cMUT, bottom). To target the construct to the correct locus by single crossover recombination, a homology region (HR) upstream of the RVP domain was used. A loxPint (red arrowhead) is positioned between the HR and the remainder of the gene, called the recodonised region (RR). Sequences encoding a 2A skip peptide and neomycin phosphotransferase (NPT) permit selection-linked integration. A second loxPint module separates the *npt* gene and a silent *gfp* gene. GFP is only expressed after RAP treatment, represented as a change in green color. For the PfDdi1_cKO line, RAP treatment results in the formation of a truncated form of PfDdi1 fused to GFP. For PfDdi1_cWT or PfDdi1_cMUT, RAP treatment results in the replacement of RR with RR2 resulting in the expression of wildtype full-length PfDdi1 or mutant D262N PfDdi1 and free GFP. The position of the primers used for the PCRs of 5’ integration and excision are shown in green and red, respectively. (**C**) Diagnostic PCRs showing integration (bottom) and excision (top) at the *PfDdi1* locus of the conditional lines. Genomic DNA was collected from the indicated lines 24h after DMSO or RAP treatment. The size of the expected PCR products is shown with arrowheads. Genomic DNA from the B11 line was used as a negative control. (**D**) WB analysis of the conditional lines. The indicated conditional lines were treated with DMSO or RAP at ring stage, and parasite lysates collected at schizont stage and analyzed by WB using either an anti-2A peptide (top) or anti-GFP (middle) antibody. Lysates from the B11 line were used as a negative control. Ponceau staining of the blot is shown to control for protein loading. (**E)** Evaluation of excision efficiency by flow cytometry. Our different lines were DMSO or RAP treated at ring stage, cultured until they reached mature schizont stage, and the GFP signal analyzed by flow cytometry after fixation of the samples. The black and green number show the percentage of GFP positive schizonts in DMSO and RAP treated parasites, respectively.

Whether *Plasmodium* Ddi1 (PfDdi1) shares conserved functions with its human and yeast homologues is unknown. Interestingly, a very recent study looked at the role of Ddi1 in the closely related apicomplexan parasite *Toxoplasma gondii* (H. Zhang et al., 2020). Although KO of TgDdi1 had no effect in parasite replication *in vitro*, its KO leads to an accumulation of ubiquitinated proteins. Importantly, TgDdi1KO parasites were less virulent *in vivo*, a phenotype than could be rescued through complementation with PfDdi1 (H. Zhang et al., 2020). In addition to the RVP domain, PfDdi1 contains an N-terminal UBL but lacks C-terminal UBA or UIM domains that can be found in ScDdi1, TgDdi1 or hDdi2. However, the UBL domain of ScDdi1 and hDdi2 have been shown to also bind ubiquitin in addition to the proteasome (Nowicka et al., 2015). Therefore, homodimeric PfDdi1 could potentially perform a proteasomal shuttle role via the dual-specificity of the UBL domain, with one binding the proteasome and the other ubiquitin. Direct attempts to KO Ddi1 in *P. berghei* have been unsuccessful (Onchieku et al., 2018), and genome-wide screens in *P. falciparum* (M. Zhang et al., 2018) and *P. berghei* (Bushell et al., 2017) found Ddi1 and multiple members of the UPS to be essential. Key differences exist between the *Plasmodium* and human UPS (Pereira et al., 2018), potentially making it an attractive drug target for malaria. In particular, the proteasome is a high priority antimalarial drug target (H. Li et al., 2016), especially as proteasome inhibitors have synergistic effects with artemisinin. Thus, understanding more about components of the *Plasmodium* UPS is a key area of research.

We hypothesised that PfDdi1 is an active aspartyl protease that is important for *P. falciparum* growth. In this study, we used a state of the art conditional genetic approach to show that PfDdi1 activity is essential for RBC invasion. Using a combination of biochemical and proteomics studies, we established a strong link between PfDdi1 and the UPS system, which we believe is conserved across eukaryotes. Importantly, we show that PfDdi1-deficient parasites are more sensitive to proteasome inhibition, thus raising the possibility that a combination of PfDdi1 and proteasome inhibitors might have synergistic effects.

## Results

### Generation of conditional truncation and conditional mutation lines

The novel conditional genetics approach used in this study utilises three technologies. The first is selection-linked integration (SLI, (Birnbaum et al., 2017)) which makes use of the viral 2A peptide to link expression of a modified *ddi1* gene with a selectable marker to rapidly select for parasites with correct integrations. In addition, the rapamycin (RAP) inducible dimerisable Cre (DiCre) system (C. R. Collins, Das, et al., 2013) was combined with artificial loxP-containing introns (loxPint) (Jones et al., 2016) to conditionally truncate or mutate the endogenous *ddi1* gene.

The conditional truncation (cKO) strategy for *PfDdi1* is outlined in Figure 1B. In the modified *ddi1* locus, *loxP-*containing introns are situated such that the entire RVP domain is excised following the activation of DiCre with RAP. At the same time, a previously silent *gfp* comes into frame with the truncated *ddi1* gene resulting in the production of a truncated protein-GFP fusion reporter.

Moreover, to determine whether any specific phenotype observed upon conditional truncation of PfDdi1 is due to its putative proteolytic activity, a RAP-inducible conditional allelic replacement strategy was devised (Figure 1B), whereby the excised RVP region is replaced with a matching region containing the catalytic Asp (wildtype, WT) or a mutant version (D262N). In these conditional allelic replacements lines, which are referred to as PfDdi1_cWT and PfDdi1_cMUT, a 2A-linked GFP is also produced following RAP treatment, acting as a reporter of excision.

These constructs were electroporated into a parasite line constitutively expressing DiCre, named B11 or 1G5 (C. R. Collins, Das, et al., 2013; Perrin et al., 2018). For all lines, integration was verified by PCR (Figure 1-figure supplement 1).

Unless otherwise indicated, treatment with DMSO or RAP was performed on synchronous ring-stage parasites. Excision of genomic DNA was demonstrated by PCR for two clones of each line (Figure 1C). Depletion of a 5 ′ integration PCR product and a reduction in the size of an excision PCR product upon RAP treatment demonstrated efficient excision in all lines. Anti-2A and anti-GFP western blot (WB) analysis on mature schizont lysates was used to assess excision at the protein level (Figure 1D). For the PfDdi1_cKO clones, RAP-mediated truncation of the C-terminal region results in the loss of the 2A peptide and formation of a truncated PfDdi1 protein fused to GFP. As expected, RAP treatment of the PfDdi1_cWT and PfDdi1_cMUT lines does not result in the loss of the 2A peptide, but in expression of free GFP (Figure 1D). Flow cytometry was also used to measure GFP fluorescence in mature schizonts and determine the excision rate (Figure 1E). After RAP treatment, ∼97% of schizonts were GFP positive for all lines, demonstrating efficient excision. PCR and WB analysis of the additional PfDdi1_cKO clones obtained in the 1G5 background are shown in Figure1- figure supplement 2.

### PfDdi1 activity is essential for parasite replication

The importance of PfDdi1 on the asexual development of *P. falciparum* was assessed using standard replication assays and plaque assays (Thomas et al., 2016). Replication assays were performed by diluting DMSO- or RAP-treated cultures to 0.1% parasitaemia and monitoring replication by flow cytometry for three cycles. Conditional truncation or mutation of PfDd1 results in a complete loss of parasite replication, while no differences between DMSO and RAP treatment were observed in our PfDdi1_cWT lines. (Figure 2A and Figure 1-figure supplement 2).

**Figure 2.**
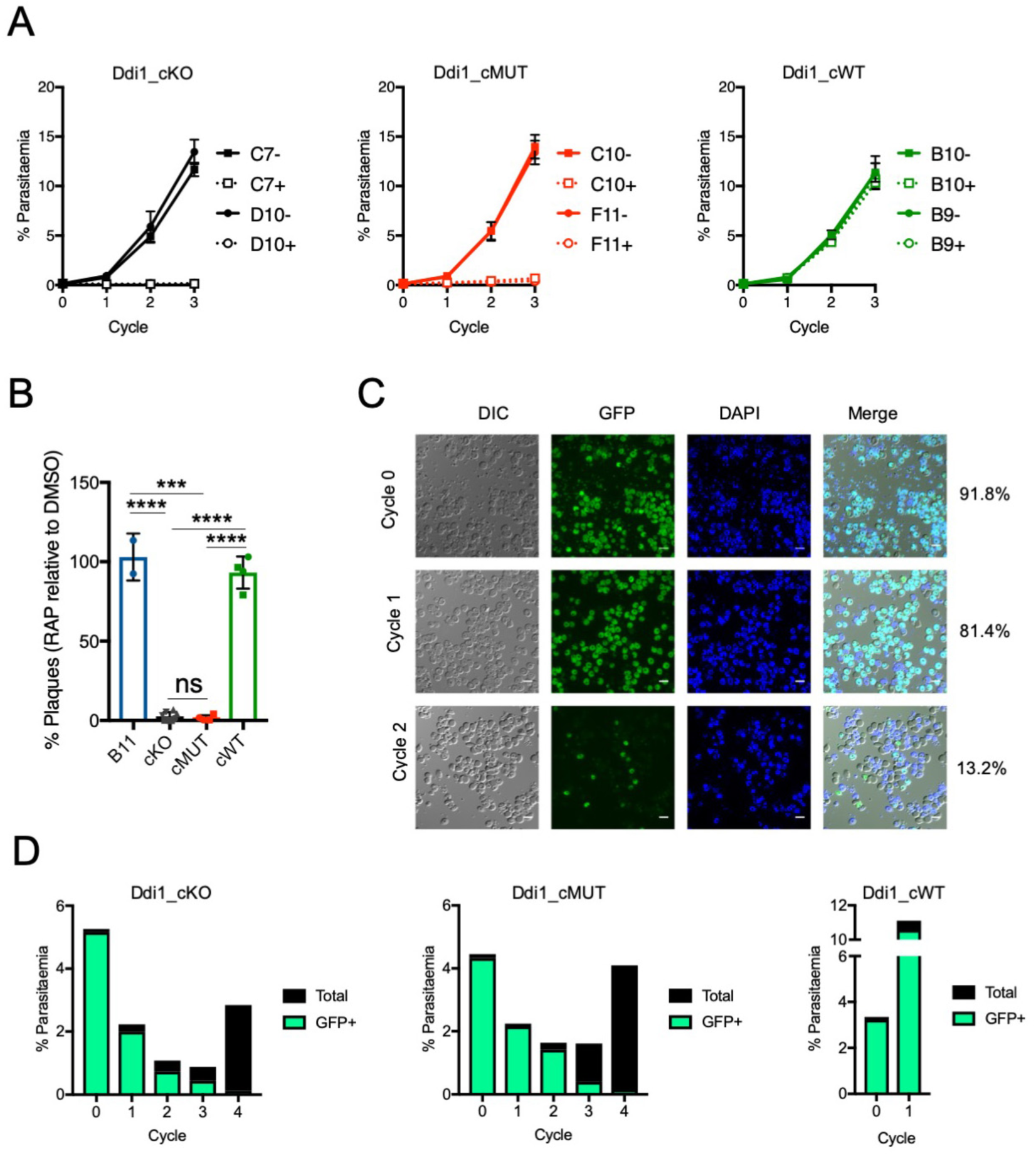
Effect of PfDdi1 disruption on parasite replication. (**A**) Replication assay. Two clones of each of our conditional lines were DMSO (-) or RAP (+) treated at ring stage, diluted to 0.1% parasitaemia, and the parasitaemia quantified over the following three cycles by flow cytometry. Error bars indicate the standard deviation of three biological replicates. (**B**) Plaque assay. After DMSO or RAP treatment, parasite cultures were diluted in flat-bottom 96-well plate at around 50 iRBCs/well. After 12 days incubation, the number of plaques were counted using an inverted microscope. For each cell line, differently shaped data points represent different clones, with two independent experiments performed for each clone. Error bars represent standard deviation (***, p<0.001; ****, p<0.0001). (**C**) Live microscopy of PfDdi1 KO parasites over 3 cycles. Synchronous PfDdi1_cKO parasites were DMSO or RAP treated and highly mature schizonts were collected at the end of the cycle of treatment (Cycle 0) and of the following two cycles (Cycles 1 and 2). These samples were stained with DAPI and analyzed by live microscopy and representative images are shown. The percentage of GFP positive schizonts at the end of each cycle is indicated on the right (n>300). No GFP positive iRBCs were observed in DMSO treated samples. (**D**) Analysis of PfDdi1-deficient parasites replication by flow cytometry over 5 cycles. Parasites at 3-5% parasitaemia were treated with RAP at ring stage and cultured for 5 cycles. At the end of each cycle samples were fixed, and the proportion of GFP positive (green) or negative (black) mature schizonts quantified by flow cytometry.

In the plaque assay (Fig 2B), parasites are plated at low parasitaemia in flat bottom 96-well plates. After 7-12 days incubation, microscopic plaques resulting from lysis of iRBCs are visible in the thin blood layer. Each plaque originates from a single iRBC, and the number of plaques is a measure of parasite replication. Loss of functional PfDdi1 upon RAP treatment (PfDdi1_cKO and PfDdi1_cMUT) results in an almost complete loss of plaque-forming ability, with any plaques remaining likely to be non-excised parasites (Figure 2B). No significant differences in plaque numbers were observed for our PfDdi1_cWT lines upon RAP treatment.

Because the DiCre system is not 100% efficient, we assessed whether the few plaques or extremely low parasitaemia observed upon RAP treatment of the PfDdi1_cKO and PfDdi1_cMUT lines were due to replication of non-excised parasites or to a very slow replication rate of PfDdi1-deficient parasites. This was assessed by monitoring GFP positive parasites after RAP treatment. The GFP fluorescence of RAP treated PfDdi1_cKO was monitored for 3 cycles by live microscopy after enrichment of mature schizonts at the end of each cycle (Figure 2C). In RAP treated samples, GFP positive schizonts were detected in all cycles, demonstrating that a subset of KO parasites was able to invade and develop in the following cycle. Even so, the 8% of non-excised parasites (GFP negative) that were present in cycle 0 quickly outcompeted the population of KO parasites (GFP positive) in culture. Indeed, by cycles 1 and 2, non-excised parasites comprised 20% and 80% of the population, respectively.

We also monitored the progression of each PfDdi1 conditional line population over 5 cycles after RAP treatment by flow cytometry. One clone of each conditional line was RAP treated, and a sample of mature schizonts fixed and analyzed at the end of each cycle. The population of excised vs non-excised parasites was quantified based on the GFP signal (Figure 2D). As previously described, the excision rate was excellent at cycle 0 (> 96%). While the PfDdi1_cWT line experienced a 3.3-fold increase in parasitaemia in the first cycle, the parasitaemia of KO and mutant cultures dropped by ∼50% each cycle, while the small proportion of non-excised parasites increased exponentially. These results indicate that PfDdi1-deficient parasites experience a profound and consistent replication defect. However, a very small proportion of merozoites (1 in the ∼40-60 merozoites that are produced per every two schizonts) are able to invade and replicate each cycle. Nonetheless, these results show that PfDdi1-deficient parasites are unable to propagate in culture, indicating that PfDdi1 is essential, and that its aspartyl protease activity is required for asexual blood stages.

### PfDdi1 is essential for RBC invasion

To determine which development stage is affected by the loss of PfDdi1 function, we used a recently developed flow cytometry assay that allows us to monitor parasite development throughout the erythrocytic cycle with hourly resolution (Bell et al., 2021). This assay combines selective DNA and RNA fluorescent dyes to measure intracellular development and changes in parasitaemia. In iRBCs, the DNA signal increases during schizogony due to DNA replication, and RNA levels increase throughout the cycle allowing us to monitor the ring to trophozoite transition. We used this assay to measure differences in parasite development of PfDdi1_cKO parasites for 80h after DMSO or RAP treatment. This approach shows that cKO of PfDdi1 has no effect in intracellular development or parasite egress, but leads to an unmistakable invasion defect (Figure 3-figure supplement 1).

To confirm these results, the invasion rate in PfDdi1_cKO, PfDdi1_cWT and PfDdi1_cMUT was assessed (Figure 3A). Mature schizonts were purified, mixed with fresh erythrocytes, and allowed to invade for 24h at 37°C. Samples were fixed before and after invasion, and the proportion of rings and schizonts determined by flow cytometry. As expected, WT allelic replacement resulted in no significant change in invasion rate. In contrast, KO or mutation of PfDdi1 led to significantly decreased invasion rates. These data demonstrate that loss of PfDdi1 results in a dramatic, but not complete, block in invasion. Interestingly, truncation of PfDdi1 had a more significant decrease in invasion rate than the D262N mutation, perhaps indicating protease-independent functions.

**Figure 3.**
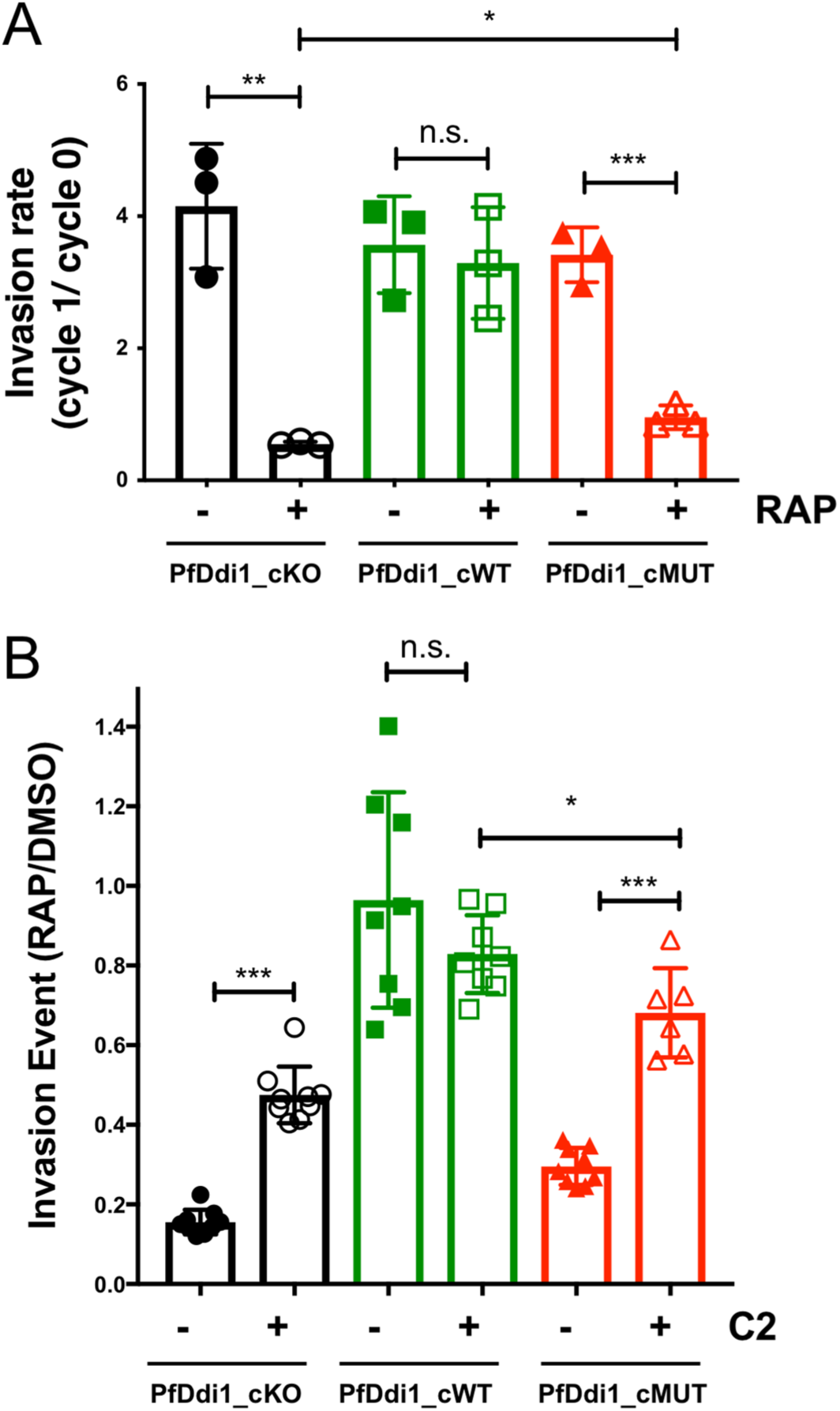
PfDdi1 is essential for RBC invasion. (**A**) Invasion assay. Schizonts from DMSO or RAP treated cultures were purified and incubated for 24h with fresh RBCs. Samples were fixed before and after invasion, and the proportions of schizonts and rings quantified by flow cytometry. Invasion rates were calculated as the number of rings formed after 24h relative to number of schizonts present at the beginning of the invasion assay. (**B**) Delaying parasite egress rescues the invasion defect. DMSO or RAP treated parasite lines were treated at 40 hpi with 1μM C2 or DMSO for 12h. Parasite cultures were then washed with media and cultured for an additional 4h before fixation and flow cytometry analysis. The bar graph shows the ratio in invasion efficiency between excised (RAP) and non-excised (DMSO) parasites as a function of C2 treatment. (**A-B**) Error bars represent standard deviation of 3 biological replicates. T-test significance values are indicated (*, p<0.05; **, p<0.01, *** p<0.001).

RBC invasion is a multistep and tightly regulated process requiring the interplay of many intracellular organelles and subcellular structures. Given the pronounced invasion defect associated with the loss of PfDdi1, we performed extensive immunofluorescence analysis (IFA) studies in mature segmented schizonts to determine whether these different components were properly formed in PfDdi1- deficient parasites. In all cases, we failed to observe differences in the localization of a variety of protein markers between DMSO and RAP treated parasites, indicating that merozoites form properly in PfDdi1-deficient parasites (Figure 3-figure supplement 2).

### The invasion defect associated with the loss of PfDdi1 function can be rescued by delaying parasite egress

We were surprised to observe that PfDdi1-deficient parasites show a pronounced invasion defect rather than a delay in intracellular parasite development. In other organisms, Ddi1 mainly localizes to the cytoplasm and its function is closely associated with the UPS. However, most proteins involved in RBC invasion are in the secretory pathway. Also, PfDdi1 is expressed as early as 24 hours post invasion (hpi), and it is therefore surprising that we only observed an invasion phenotype. Finally, if PfDdi1 is involved in protein quality control, we might have expected to see a delay in schizogony.

To address this point, we compared the invasion efficiency of our different conditional lines depending on whether or not egress had been stalled with the cGMP dependent kinase (PKG) inhibitor Compound 2 (C2). The rationale behind this experiment was to test whether the invasion defect could be rescued if parasites were given more time to mature before egress. Prior to egress, cytosolic levels of cGMP increase, leading to activation of PKG. This triggers the release of micronemal and exonemal proteins that are important for egress and invasion (C. R. Collins, Hackett, et al., 2013). Treatment with C2 arrests parasites 15 min before egress, a process that is reversible if the inhibitor is later washed away from the cultures.

After DMSO or RAP treatment of the different conditional lines at ring stage, cultures were treated either with C2 or DMSO at 40 hpi. After 12h (52 hpi), C2 or DMSO were washed out, and parasites allowed to egress and invade for an additional 4h before fixing them for flow cytometry analysis. No significant difference in the proportion of schizonts that had egressed between DMSO and RAP treatment was observed in any of the lines. However, in PfDdi1-deficient parasites (PfDdi1_cKO and PfDdi1_cMUT) we observed a very significant recovery of invasion efficiency in C2 treated parasites (Figure 3B). No differences were observed for the PfDdi1_cWT control line. These results point to PfDdi1-deficient parasites having a defect in merozoite maturation that can be resolved if parasites are allowed more time to mature before egress. Note that conditional truncation of PfDdi1 had a more significant effect in RBC invasion than mutation of its catalytic residue, supporting the idea that PfDdi1 may have both catalytic and non-catalytic functions.

### PfDdi1-deficient parasites show a delay in cytokinesis

To further explore this potential delay in merozoite maturation, PfDdi1_cMUT parasites were DMSO or RAP treated, and schizonts purified at 44 hpi and cultured until they reached mature schizonts (48 hpi) in the presence of C2. Samples were analyzed by serial block face scanning electron microscopy (SBF_SEM), which provides 3D structural information of the cell. Using this method, we could not observe any obvious defect in the overall structure of merozoites. However, quantification of the proportion of segmented vs non-segmented schizonts reveals a higher proportion of segmented schizonts in the DMSO control compared to the RAP treated population (62% versus 38%) (Figure 4A). Analysis of the number of nuclei in these EM images showed no overall significant difference between DMSO and RAP treated parasites. These results clearly show that loss of PfDdi1 activity has no effect in nuclear division but results in a delay in cytokinesis. Videos showing each SBF-SEM section used for quantification are shown in Figure 4-figure supplement 1-7.

**Figure 4.**
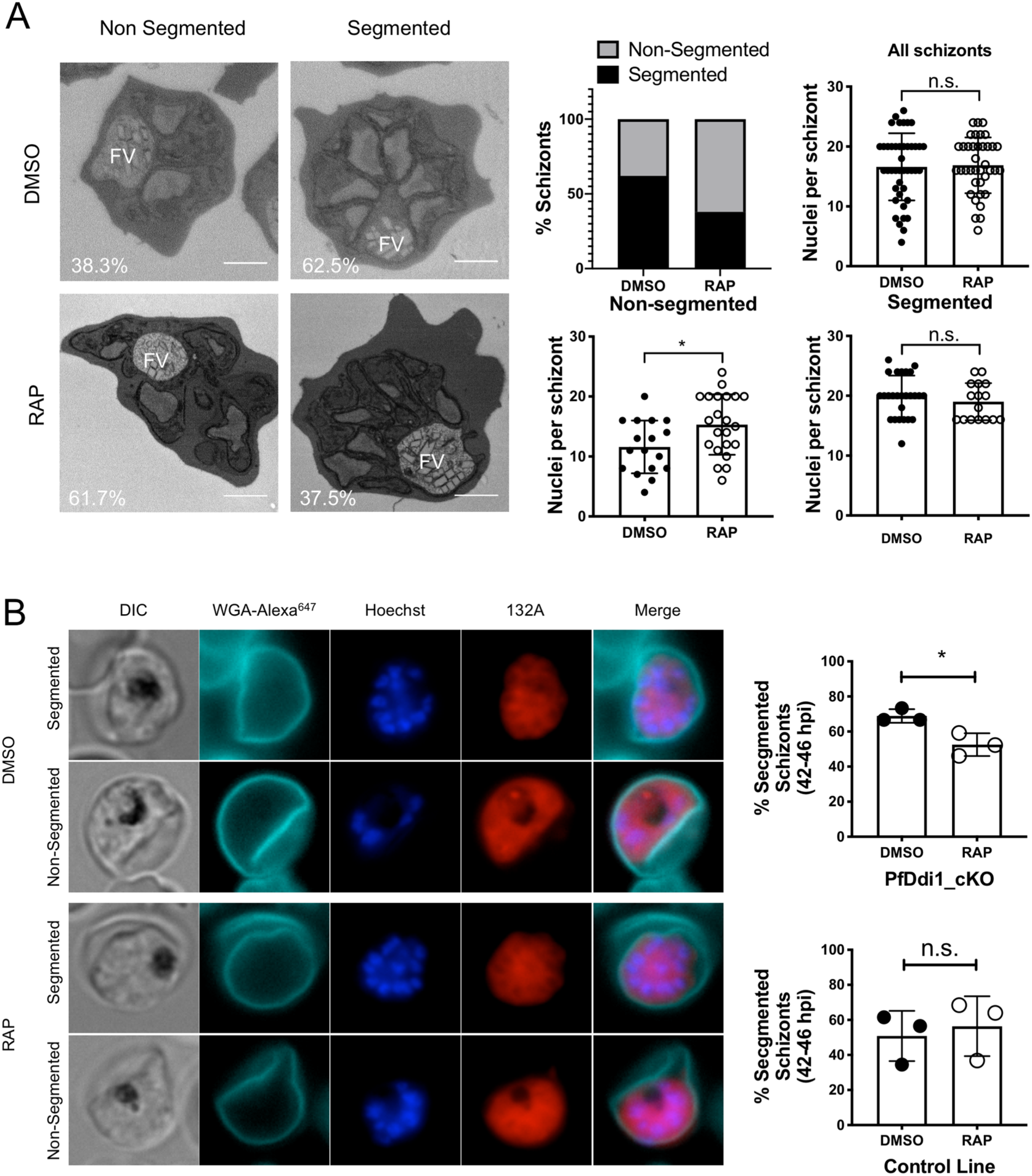
Disruption of PfDdi1 results in a delay in merozoite maturation. (**A**) Quantification of segmented vs non-segmented schizonts by EM. Representative images of segmented and non-segmented schizonts obtained by SBF_SEM for the PfDdi1_cMUT line upon DMSO or RAP treatment. The left graph illustrates that a larger proportion of non-segmented schizonts were present in the RAP treated culture. The number of nuclei per schizont in either segmented or non-segmented schizonts are shown in the middle and right graphs, respectively. (**B**) Quantification of segmented vs non-segmented schizonts by fluorescence microscopy. Fixed samples collected at 42, 44, and 46 hpi from the PfDdi1_cKO^1G5^ and 1G5, were used to look at differences in the morphology of the schizonts by fluorescence microscopy between DMSO and RAP treated samples. DNA and RNA were stained with Hoechst and the 132A dye, respectively. In addition, the RBC surface was stained with Alexa647-conjugated WGA. Samples were visualized in an inverted microscope, and 20-60 schizonts were counted per sample. Representative images of segmented and non-segmented schizonts are shown for DMSO and RAP treated parasites. In segmented schizonts staining with the 132A RNA dye clearly shows a bundle of grape pattern representative of individual merozoites having been formed. In non-segmented schizonts the RNA staining is continuous and occupies most of the iRBC area. The bar graph shows that RAP treated parasites have a significantly higher number of non-segmented schizonts. Each data point corresponds to a different time point. (**A-C**) Error bars represent standard deviation. T-test significant values are shown (*, p<0.05; **, p<0.01, *** p<0.001).

To confirm this delay in cytokinesis, we look at the morphology of DMSO or RAP treated PfDdi1_cKO parasites a few hours before egress (42, 44 and 46 hpi) by fluorescent microscopy using the DNA (Hoechst) and RNA (132A) dyes described above (Figure 3-figure supplement 1). The RNA dye stains the parasite cytosol and nucleus. At the end of schizogony, segmented vs non-segmented schizonts can be visualized depending on the pattern of RNA staining. In non-segmented schizonts, the RNA signal is evenly distributed and shows a single circular staining occupying most of the iRBCs. In segmented schizonts, the delineation of individual merozoites can be visualized as a bundle of grapes shape in the RNA channel (Bell et al., 2021). Samples were also stained with wheat germ agglutinin conjugated to AlexaFluor647 (WGA-647), which by to lectins on the surface of the RBC (Figure 4B).

Quantification of the number of segmented vs non-segmented schizonts in this samples also indicate a higher proportion of segmented schizonts in DMSO treated parasites. Overall, these results suggest that PfDdi1-deficient parasites show a small delay in merozoite formation at the end of schizogony. This might result in a smaller number of mature merozoites being released at the time of egress and could explain the strong invasion defect.

### PfDdi1 localizes to puncta in the cytoplasm of *P. falciparum*

Previous attempts by our group and others to directly tag *Plasmodium* Ddi1 with a triple HA tag or a fluorescent protein have been unsuccessful. However, in our conditional lines, PfDdi1 is expressed with a C-terminal 2A peptide showing that small C-terminal tags are tolerated. To explore the localisation and interactome of PfDdi1, a new cell line was generated (PfDdi1_cHA) to conditionally introduce a single HA tag at the C-terminal. The PfDdi1_cWT construct was modified by incorporating an HA tag at the 3’ end of the second recodonised region instead of 2A-linked *gfp*. This construct was electroporated into the B11 line. In the resultant PfDdi1_cHA lines, PfDdi1-2A is produced prior to excision, and PfDdi1-HA after RAP treatment. Validation of this line and confirmation that the HA tag did not impact parasite replication is shown in Figure 5-Supplement 1.

The localization of PfDdi1-HA was determined at 30, 42, and 48 hpi by confocal microscopy (Figure 5A). At each time point, PfDdi1-HA presented as puncta within the parasite. At 30 hpi it is particularly clear that these puncta are spread throughout the cytoplasm and excluded from the food vacuole. In later stages, PfDdi1-HA appears to be absent from the nuclei. PfDdi1-HA labelling was compared to that of the surface protein MSP1 and of the rhoptry protein RhopH2, as a marker of the apical end of merozoites (Figure 5B). These IFAs show that PfDdi1-HA is present within the confines of MSP1 staining, and that the foci do not co-localize with RhopH2. In addition, there appears to be multiple puncta distributed throughout each daughter merozoite indicating that PfDdi1 has a cytoplasmic, non-apical, localization.

**Figure 5.**
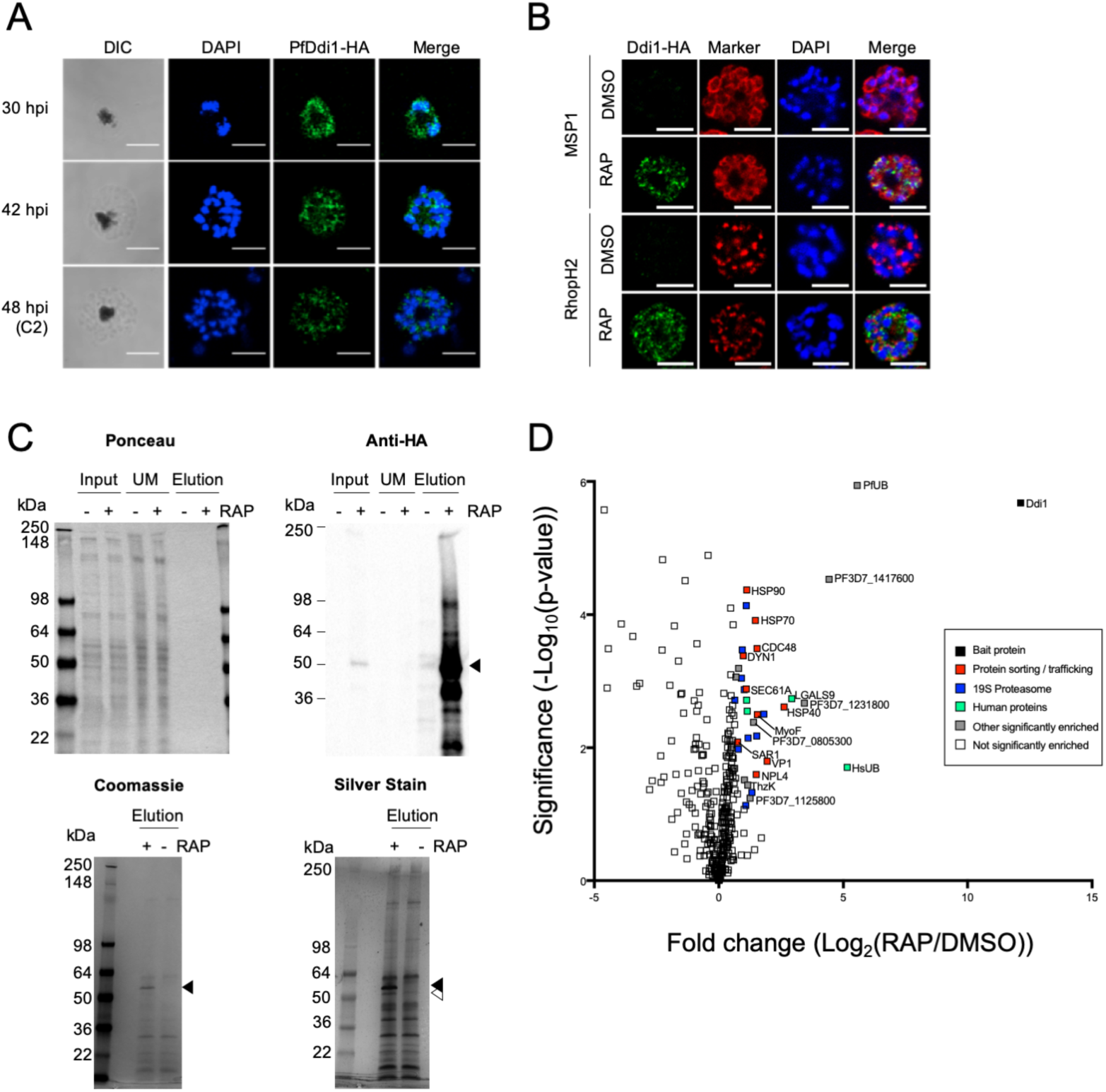
PfDdi1 localizes to the cytosol and interacts with components of the UPS. (**A**) Localization of PfDdi1-HA. Parasite expressing PfDdi1-HA were collected at early (30 hpi), mid (42 hpi) and late (48 hpi) schizont stages. IFA shows punctate cytosolic localization of PfDdi1-HA. Note that PfDdi1-HA does not colocalize with the nuclei or the food vacuole, where the hemozoin crystal resides and can be visualized in the DIC channel. (**B**) IFA analysis of mature schizonts after DMSO or RAP treatment of the PfDdi1_cHA line. Colocalization with merozoite surface (MSP1) and rhoptry (RhopH2) markers clearly show that PfDdi1-HA resides within merozoites and that it does not localize to the apical end. (**C**) Immunoprecipitation of PfDdi1. Parasite lysates of DMSO (-) or RAP (+) treated PfDdi1_cHA were collected at mature schizont stage, incubated with streptavidin beads, and the pull-down material eluted by boiling the beads with loading buffer. Input, unbound (UM) and elution material were analyzed by WB. Ponceau staining was used to control for protein loading. Elution samples were stained with coomassie and silver stain. The black and white arrowheads indicate the band corresponding to PfDdi1-HA and a potential interacting protein, respectively. (**D**) Proteomic analysis of PfDdi1-HA pull-down. Volcano plot shows the average log(2) fold enrichment of proteins between DMSO and RAP treated PfDdi1_cHA parasites as a function of the significance value based on three technical replicates. Significantly enriched proteins are shown in different colors depending on their predicted biological function as indicated in the legend.

### Pull-down of PfDdi1-HA interacting partners identifies proteins implicated in protein sorting and degradation

The conditional tagging of PfDdi1 allows pull-down of this protein from parasite material along with any interacting partners. Using anti-HA beads, PfDdi1-HA was purified from the soluble lysate of PfDdi1_cHA excised schizonts, alongside pulldowns from the non-excised controls. Protein was eluted by boiling in SDS-containing buffer and the purification was validated by WB (Figure 5C).

The immunoprecipitation eluates were analyzed by liquid chromatography tandem mass spectrometry (LC-MS/MS). A volcano plot of the fold difference vs significance between DMSO and RAP treated samples is shown in Figure 5D. The significantly enriched proteins are listed in Table 1. As anticipated, PfDdi1 was the most enriched protein (4500-fold) in the pull-down, followed by ubiquitin. Remarkably, 11 putative subunits of the proteasome 19S regulatory particle (RP) were significantly enriched, including 5 of the 6 AAA ATPases present in the 19S base, and 5 of the 9 proteins that form the lid of the 19S RP (Aminake et al., 2012). Another significantly enriched protein was CDC48, a ubiquin-directed unfoldase/segregase implicated in many cellular processes, including the translocation of misfolded proteins from the ER in concert with NPL4 and SEC61A, both of which were also enriched in the pull-down. Three heat shock proteins, HSP90, HSP70-1 and an HSP40 (DnaJ) were also significantly enriched as well as several proteins involved in trafficking, i.e. V-type H(+)- translocating pyrophosphatase (VP1), GTP binding protein Sar1, myosin F (MyoF), and dynamin-like protein 1 (DYN1). Overall, many of the proteins that co-purified with PfDdi1 likely have roles in the UPS and protein quality control in *P. falciparum*. If PfDdi1 directly binds both ubiquitin and the proteasome, then it may be capable of performing a proteasome shuttle function, as described for yeast Ddi1 (Ivantsiv et al., 2006; Kaplun et al., 2005).

**Table 1.** Significantly Enriched Proteins following the PfDdi1-HA Pull-down

| Gene IDs | Name | Description | Fold change |
| --- | --- | --- | --- |
| PF3D7_1409300 | Ddi1 | DNA damage-inducible protein 1, putative | 4508 |
| PF3D7_1365900;<br>PF3D7_1211800 | PfUB | Ubiquitin | 47 |
| PF3D7_1417600 |  | conserved Plasmodium protein, unknown function | 22 |
| PF3D7_1231800 |  | asparagine-rich protein, putative | 11 |
| PF3D7_0823800 | HSP40 /<br>DnaJ | DnaJ protein, putative | 6.26 |
| PF3D7_1456800 | VP1 | V-type H(+)-translocating pyrophosphatase, putative | 3.4 |
| PF3D7_1129200 | RPN7 | 26S proteasome regulatory subunit RPN7, putative | 3.5 |
| PF3D7_0805300 |  | conserved Plasmodium protein, unknown function | 2.9 |
| PF3D7_0619400 | CDC48 | cell division cycle protein 48 homologue, putative | 2.9 |
| PF3D7_1248900 | RPT6 | 26S protease regulatory subunit 8, putative | 2.9 |
| PF3D7_0507700 | NPL4 | nuclear protein localization protein 4, putative | 2.8 |
| PF3D7_0818900 | HSP70 | heat shock protein 70 | 2.8 |
| PF3D7_1329100 | MyoF | myosin F, putative | 2.6 |
| PF3D7_1466300 | RPN2 | 26S proteasome regulatory subunit RPN2, putative | 2.5 |
| PF3D7_1125800 |  | kelch domain-containing protein, putative | 2.4 |
| PF3D7_1130400 | RPT5 | 26S protease regulatory subunit 6A, putative | 2.3 |
| PF3D7_1239600 | ThzK | hydroxyethylthiazole kinase | 2.2 |
| PF3D7_0708400 | HSP90 | heat shock protein 90 | 2.2 |
| PF3D7_1338100 | RPN3 | 26S proteasome regulatory subunit RPN3, putative | 2.2 |
| PF3D7_1346100 | SEC61A | protein transport protein SEC61 subunit alpha | 2.1 |
| PF3D7_1017900 | RPN5 | 26S proteasome regulatory subunit p55, putative | 2.1 |
| PF3D7_1027300 | nPrx | peroxiredoxin | 2.0 |
| PF3D7_0912900 | RPN8 | 26S proteasome regulatory subunit RPN8,<br>putative | 2.0 |
| PF3D7_1145400 | DYN1 | dynammin-like protein | 2.0 |
| PF3D7_1311500 | RPT1 | 26S protease regulatory subunit 7, putative | 1.9 |
| PF3D7_1008400 | RPT2 | 26S protease regulatory subunit 4, putative | 1.9 |
| PF3D7_0519400 | RPS24 | 40S ribosomal protein S24 | 1.7 |
| PF3D7_0413600 | RPT3 | 26S protease regulatory subunit 6B, putative | 1.7 |
| PF3D7_0416800 | SAR1 | small GTP-binding protein sar1 | 1.7 |
| PF3D7_1105400 | RPS4 | 40S ribosomal protein S4, putative | 1.6 |
| PF3D7_1368100 | RPN11 | 26S proteasome regulatory subunit RPN11,<br>putative | 1.6 |

### PfDdi1-deficient parasites are more sensitive to proteasome inhibition

Given the association between PfDdi1 and the proteasome, we tested the ability of PfDdi1-deficient parasites to recover from proteasome inhibition. After DMSO or RAP treatment of PfDdi1_cKO, PfDdi1_cWT and PfDdi1_cMUT, parasites were treated for 4h (starting at 36 hpi) with different concentrations of the proteasome inhibitor bortezomib (BTZ). After washing away the inhibitor, the parasite cultures were allowed to egress and invade for an additional 24h. After fixation, parasitaemia was quantified by flow cytometry and fitted to a dose response curve. Strikingly, a more than 10-fold decrease in the EC_50_ value was observed upon RAP treatment of PfDdi1_cKO (Figure 6A), demonstrating that parasites lacking PfDdi1 are much more sensitive to proteasome inhibition than controls. Surprisingly, conditional mutation of the catalytic Asp has no effect on the sensitivity of parasites to BTZ treatment. This result again highlights that PfDdi1 has both catalytic and non-catalytic functions.

**Figure 6.**
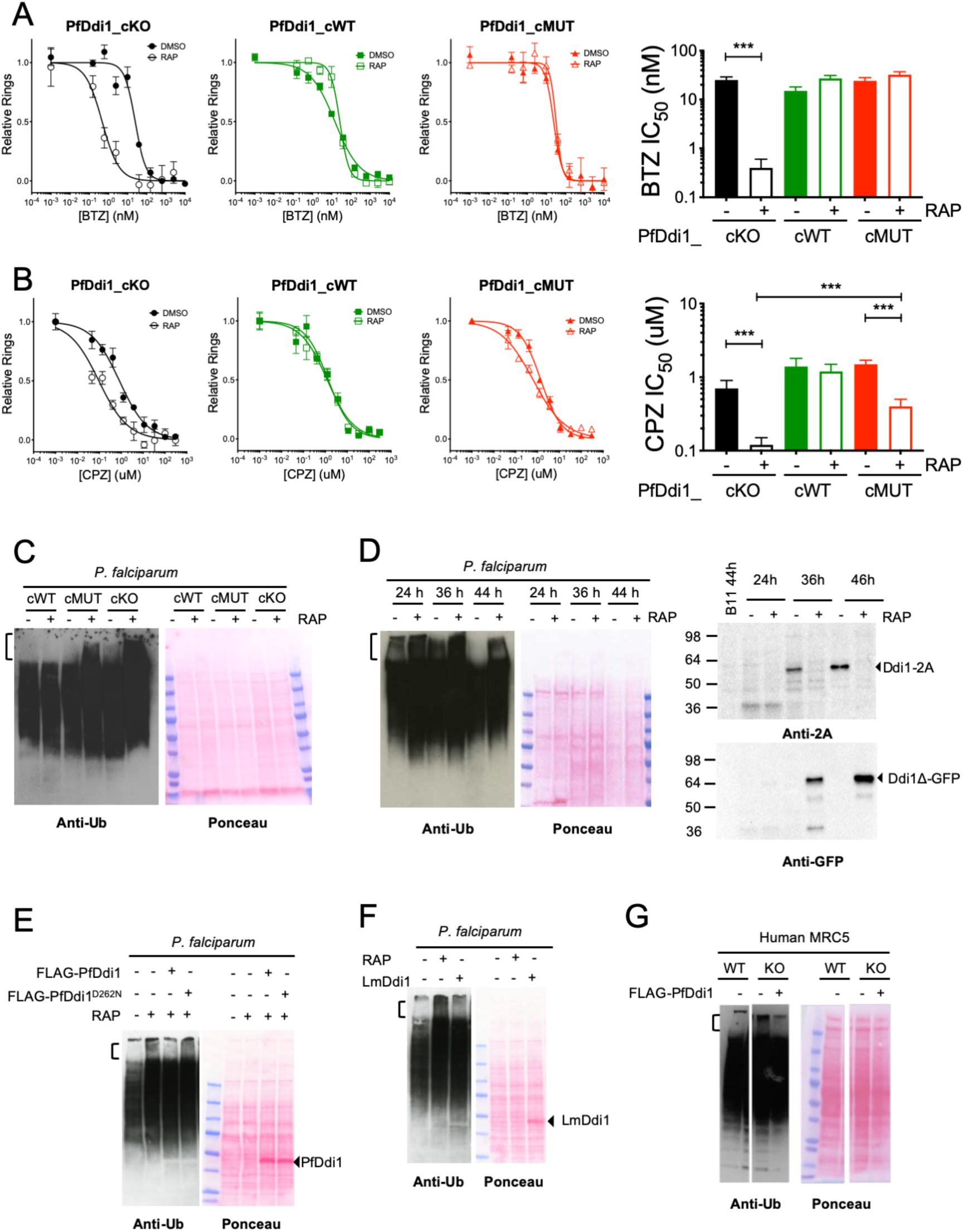
PfDdi1-deficient parasites are more sensitive to proteasome inhibition, and PfDdi1 cleaves highly polyubiquitinated proteins. (**A**) DMSO or RAP treated conditional lines were treated for 4h with different concentrations of BTZ. Parasites were cultured for an additional 24h, and parasitaemia quantifed by flow cytometry analysis. (**B**) Parasite cultures were treated for 24h with a dose response of CPZ starting at 36 hpi. Samples were then fixed and analyzed by flow cytometry. (**A-B**) Parasitaemia was fitted to a dose response curve to obtain IC_50_ values (bar graphs). Error bars represent the standard deviation of three biological replicates. T-test significant values are also shown (*, p<0.05; **, p<0.01, *** p<0.001). (**C**) Accumulation of HMWUPs in PfDdi1-deficient parasites. Schizont lysates obtained after DMSO or RAP treatment of our conditional lines were analyzed by anti-ubiquitin WB. (**D**) The accumulation of HMWUPs is inversely correlated with PfDdi1 expression. Parasite lysates from the PfDdi1_cKO line were collected at 24, 36, and 46 hpi after DMSO or RAP treatment at ring stage. WB analysis using anti-2A and anti-GFP antibodies shows efficient excision upon RAP treatment. Anti-ubiquitin WB shows an accumulation of HMWUPs upon RAP treatment. (**E**) PfDdi1 cleaves *Plasmodium* HMWUPs. PfDdi1 KO parasite lysates were treated either with FLAG-PfDdi1 or FLAG-PfDdi1^D262N^, and cleavage of HMWUPs analysed by WB. (**F**) LmDdi1 cleaves *Plasmodium* HMWUPs. PfDdi1 KO schizont lysates were treated with or without recombinant LmDdi1 and analyzed by anti-ubiquitin WB. (**G**) *Plasmodium* PfDdi1 cleaves human HMWUPs. Lysates from hDdi2 KO MRC5 cells were treated with or without FLAG-PfDdi1 and analyzed by anti-ubiquitin WB. (**C-G**) Ponceau staining is shown for all WBs to control for protein loading, the area of the blot corresponding to HMWUPs is show with a braket symbol.

The proteasome is activated and regulated by a variety of regulatory proteins and protein complexes with which it is in continuous equilibrium. The two most prominent forms of the proteasome found in the cell are the core 20S form, involved in the degradation of cytosolic proteins, and the capped 26S form, which is responsible for the degradation of ubiquitinated proteins and the one involved in the ER-associated degradation (ERAD) pathway. Capzimin (CPZ) is a natural product that has been recently identified as an inhibitor of the ATPase activity of the Rpn11 subunit of the proteasome RP (J. Li et al., 2017). This activity is necessary for the ATP-dependent unfolding of ubiquitinated proteins targeted for proteasomal degradation. CPZ should therefore allow us to discriminate between the function of the 26S proteasome compared to the overall function of the proteasome activity in its different forms (inhibition by BTZ). Our different lines were treated at 36 hpi with different concentrations of CPZ for 24h, and parasitaemia quantified by flow cytometry. In this case, both PfDdi1_cKO and PfDdi1_cMUT parasites were more sensitive to CPZ inhibition after RAP treatment although the effect was more pronounced on the PfDdi1_cKO line (Figure 6B). No significant differences were observed for the PfDdi1_cWT line.

The differences observed in the sensitivity of PfDdi1 KO and MUT parasites to BTZ and CPZ treatment is intriguing but could be explain by the fact that the proteaseome in involved in a variety of biological functions and that PfDdi1 has both catalytic and non-catalytic functions. In the context of BTZ treatment, all the functions of the proteasome are inhibited. Therefore, parasite death occurs due to inhibition of the different biological pathways that are regulated by the proteasome. In this context, the loss of PfDdi1 activity might not have an additional impact on parasite replication. However, because conditional truncation of PfDdi1 affects both its catalytic and non-catalytic functions, this likely translates into a greater sensitivity to BTZ. On the other hand, CPZ only affects the 26S proteasome functions and not those of the 20S proteasome. In this context, the loss of PfDdi1 activity might be sufficient to observe increased sensitivity to 26S proteasome inhibition. Overall, these results demonstrate that loss of PfDdi1 sensitises parasites to proteasome inhibition, and alongside the PfDdi1 interactome data, strongly suggest that PfDdi1 performs important functions within the *P. falciparum* UPS.

### Ddi1-dependent processing of ubiquitinated proteins

In yeast and human MRC-5 cells, loss of ScDdi1 (Saeki et al., 2002) or hDdi2 (Dirac-Svejstrup et al., 2020) results in the accumulation of a subset of high molecular weight ubiquitinated proteins (HMWUPs). This effect also occurred in MRC-5 cells expressing a catalytically inactive hDdi2 mutant. Crucially, addition of recombinant hDdi2 to hDdi2 KO lysates caused the processing of these unknown HMWUPs, providing the first evidence that hDdi2 has proteolytic activity. Importantly, the processing/cleavage of these HMWUPs by recombinant hDdi2 is independent of proteasome activity (Dirac-Svejstrup et al., 2020).

We now explored whether PfDdi1 displays a similar activity. Mature schizont samples from the PfDdi1_cKO, PfDdi1_cWT and PfDdi1_cMUT lines were subjected to anti-ubiquitin WB (Figure 6C). In both PfDdi1 KO and MUT samples, a smear at the top of the blot indicated the accumulation of HMWUPs, strikingly similar to what was observed in human cells (Dirac-Svejstrup et al., 2020). In contrast, in the DMSO controls and the WT allelic replacement line, this region of the blot was clear. This experiment indicates that the proteolytic activity of PfDdi1 prevents the accumulation of these unknown ubiquitinated proteins. Importantly, anti-ubiquitin WB of Ddi1_cKO lysates collected at 24, 36 and 44 hpi suggest that such HMWUPs are present naturally at 24 hpi rather than being only an artefact of PfDdi1 loss (Figure 6D). Indeed, analysis of PfDdi1-2A expression shows that the presence of HMWUPs is inversely correlated with PfDdi1 levels during the erythrocytic cycle (Figure 6D).

### PfDdi1 cleaves HMWUPs

We recently showed that hDdi2, but not its catalytically dead counterpart, is able to cleave HMWUPs present in extracts from hDdi2 KO human cells (Dirac-Svejstrup et al., 2020), strongly suggesting that such proteins are the natural substrates of hDdi2 activity. To test whether PfDdi1 displays a similar activity, we recombinantly expressed and purified full length PfDdi1 and the D262N inactive mutant, both containing an N-terminal FLAG tag (FLAG-PfDdi1 and FLAG-PfDdi1^D262N^ (Figure 6-figure supplement 1). Gratifyingly, recombinant FLAG-PfDdi1, but not FLAG- PfDdi1^D262N^, was able to cleave HMWUPs present in PfDdi1 KO lysates (Figure 6E). Remarkably, we also found that recombinant leishmania LmDdi1 is able to cleave HMWUPS present in PfDdi1 KO lysates (Figure 6F), and that FLAG-PfDdi1 cleaves HMWUPs present in human MRC5 hDdi2 KO cell lysates (Figure 6G). Overall, these results show that PfDdi1 has proteolytic activity directed towards a variety of proteins with long ubiquitin chains and that its biochemical function is conserved across eukaryotes.

## Discussion

In this study, we identify Ddi1 as an essential aspartyl protease in the replication of *P. falciparum* asexual blood stage parasites. Loss of PfDdi1 and its activity results in a profound invasion defect. PfDdi1 is cytoplasmic and interacts with multiple components of the UPS. Indeed, we find that its loss sensitizes *P. falciparum* blood-stage parasites to proteasomal inhibition and that PfDdi1 is required for the turnover of hyperubiquitinated proteins.

### The proteolytic activity of PfDdi1 is critical for erythrocyte invasion

PfDdi1 has an essential function in the replication of the malaria parasite during the erythrocytic cycle. Disruption of PfDdi1, either through truncation or point-mutation of its catalytic site, results in a pronounced invasion defect. Such a specific invasion defect phenotype was surprising for several reasons. First, PfDdi1 starts to be expressed at trophozoite stage but its disruption has no effect until 24h later. Second, because Ddi1 has been associated with protein quality control and DNA replication control in other organisms, we might have expected PfDdi1 disruption to have an effect during schizogony. And third, most proteins involved in invasion are in the secretory pathway but our PfDdi1-HA localization studies show that PfDdi1 is primarily a cytosolic protein. Although Ddi1 has been shown to be involved in protein trafficking in other organisms (Kama et al., 2018; Lustgarten & Gerst, 1999), our IFA studies in mature schizonts indicate that disruption of PfDdi1 does not appear to result in mislocalization of proteins involved in invasion.

RBC invasion is a very inefficient process requiring the timely coordination of many biological processes and the maturation of a variety of proteins. RBC invasion is the bottleneck of the erythrocytic cycle, and under optimal *in vitro* conditions RBC invasion efficiency is usually only in the order of 20-30%. In addition, extracellular merozoites are only viable for a few minutes before they lose their ability to invade RBCs. Therefore, if merozoite maturation is impacted then a reduction in invasion efficiency would be expected. Indeed, stalling egress with a PKG inhibitor, thus allowing more time for merozoites to mature, results in a very significant recovery of the invasion defect in PfDdi1-deficient parasites. In addition, a more detailed analysis of the phenotype associated with the loss of PfDdi1 function by EM and fluorescence microscopy suggests a delay in cytokinesis, which may contribute to or explain the poor invasion efficiency of Ddi1-deficient merozoites.

### Support for a proteasome shuttle function for PfDdi1

We find that PfDdi1-HA is primarily present in the cytoplasm of the parasite and exhibits a non-uniform punctate localization in schizonts and multiple foci per merozoite after cytokinesis. This localization might represent an association with proteasomes, which have been found to have a largely punctate cytoplasmic localization in *P. falciparum* (Aminake et al., 2012). Consistent with this idea, a remarkable 11/19 proteins of the 19S RP were significantly enriched in the PfDdi1-HA pull-down, supporting the idea that PfDdi1 interacts directly with the proteasome, possibly as shuttle, as has been demonstrated for ScDdi1 and hDdi2 (G. A. Collins & Goldberg, 2020; Ivantsiv et al., 2006; Kaplun et al., 2005; Voloshin et al., 2012).

Further evidence of the functional connection between PfDdi1 and the UPS comes from our proteasome inhibition studies. We show that conditional truncation of PfDdi1 results in higher sensitivity to BTZ, an inhibitor of all the proteolytic subunits of the proteasome, and to CPZ, an inhibitor that has been shown to block the ATPase activity of human Rpn11 required for the unfolding of ubiquitinated proteins. Interestingly and somewhat surprisingly, mutation of PfDdi1’s active site results in increased sensitivity to CPZ, but not to BTZ. These differences likely illustrate the presence of both a catalytic and non-catalytic functions for PfDdi1, and the fact that the proteasome is involved in innumerable biological functions. A role for PfDdi1 beyond its proteolytic activity is also evident from the fact that cKO of PfDdi1 has a stronger effect on invasion efficiency than its conditional point-mutation. Mutation of PfDdi1 results in inhibition of its proteolytic activity, but its truncation also results in the loss of the dimerization domain, which may prevent PfDdi1 from performing its putative proteasome shuttling function.

### A conserved biochemical function for Ddi1 proteins

While the presence or absence of UBL or UBA domains varies between different eukaryotic Ddi1 proteins, the RVP domain remains well conserved, suggesting a conserved catalytic role. As in budding yeast (Yip et al., 2020), humans cells (Dirac-Svejstrup et al., 2020) and *Toxoplasma gondii* (H. Zhang et al., 2020), we find that loss of PfDdi1 results in the accumulation of ubiquitylated proteins, which are of high molecular weight, almost certainly indicating that they contain very long ubiquitin chains (Dirac-Svejstrup et al., 2020; Yip et al., 2020). Importantly, parasites that are reliant on mutant PfDdi1 also show a build-up of these hyper-ubiquitinated species. Furthermore, incubation of PfDdi1 KO lysates with recombinant PfDdi1 results in the cleavage of these HMWUPs, while no processing was observed upon addition of inactive PfDdi1. Thus, we provide strong evidence to support the idea that, as in human cells, PfDdi1 is responsible for the direct cleavage of these hyper-ubiquitinated species. Our time course analysis suggests that such proteins are naturally formed during the parasite lifecycle, rather than being an artifact due to the loss of PfDdi1, and that they are being processed once PfDdi1 is expressed in the second half of the erythrocytic cycle. Finally, we clearly show that the biochemical function of PfDdi1 is conserved across eukaryotes. Indeed, addition of recombinant PfDdi1 to human hDdi2 KO lysates also results in potent cleavage of human hyper-ubiquitinated proteins, and the same result was observed with the addition of *leishmania* LmDdi1 to PfDdi1 KO lysate. This result is especially relevant given that mammals (*Homo sapiens*), apicomplexan (*Plasmodium*), and kinetoplasts (*Leishmania*) diverged at a very early point during evolution. Although other possibilities cannot be ruled out, these results appear to support the idea that Ddi proteins digest ubiquitin chains, rather than merely site-specifically in the target protein as in the case of NRF1 (Dirac-Svejstrup et al., 2020).

### PfDdi1 as a novel antimalarial target

The essentiality of PfDdi1 and the importance of its proteolytic activity provides an important, initial validation of PfDdi1 as an antimalarial target. Understanding the importance of PfDdi1 throughout the life cycle, particularly the liver and sexual blood stages, is a crucial next step. In the rodent malaria parasite *P.* berghei, Ddi1 was classified as “likely essential” in a genome-wide screen (Bushell et al., 2017; Schwach et al., 2015), raising hope that Ddi1 is essential across the *Plasmodium* genus, and thus, also represents a good drug target in distantly related human infective species, such as *P. vivax*. Targeting PfDdi1 could have synergistic effects with the inhibition of other antimalarial targets involved in the UPS response or drugs that activate the UPS in response to cellular stress. This is exemplified here by the higher sensitivity of PfDdi1-deficient parasites to proteasome inhibition.

The main challenge to protease drug discovery is achieving target specificity as proteases of the same families have conserved mechanism of actions. The RVP domain is rare and may mean that Ddi1 can be targeted over other aspartyl proteases. Indeed, HIV proteñase, a close homologue of Ddi1, is the target of a multitude of highly specific drugs (Ghosh et al., 2016). If PfDdi1 has ubiquitin-dependent activity then this novel mechanism may make it possible to target it in a highly specific manner. Although hDdi2 can be deleted in human cells (Dirac-Svejstrup et al., 2020; Koizumi et al., 2016; Kottemann et al., 2018), recent studies in mouse have shown that Ddi2 is essential for embryonic development and its KO induces proteotoxic stress and proteasome impairment (Siva et al., 2020). In any case, given that effective malaria therapies only require very short treatment times (1-3 days), crossreactivity of PfDdi1 inhibitors with hDdi2 might not be a problem. Further work to mechanistically understand Ddi1 activity and identify specific inhibitors are critical to explore this protein as a drug target in malaria and cancer.

## Materials and methods

### Generation of constructs for conditional modification of *PfDdi1*

All primers used for PCR in this study are listed in Table supplement 1. The *PfDdi1* homology region (HR) was amplified from *P. falciparum* 3D7 genomic DNA while synthetic DNA (GeneArt) was ordered containing the final 40 nucleotides of the PfDdi1 HR, a loxPint module, and the RR (recodonised to *E. coli*) of *PfDdi1*, which excluded the stop codon. The *PfDdi1* HR and synthetic DNA PCR were cloned into BglII and SalI linearized pT2A_FIKK10-cKO vector (kind gift of Dr Moritz Treeck, The Francis Crick Institute) (Davies et al., 2020) in an In-Fusion (ClonTech) reaction to produce the plasmid pT2A_Ddi1_cKO. The construct was used to transform XL10-Gold Ultracompetant cells (Agilent).

To generate the pT2A_PfDdi1_cWT construct, a RR2 (recodonised to *P. falciparum*) was ordered as a gBlock gene fragment (Integrated DNA Technologies) with a SalI site and 2A peptide sequence at the 3’ end. The end of the *npt* gene and loxPint module were PCR-amplified from AvaI linearized pT2A_Ddi1_cKO and subsequently combined with the RR2 gBLOCK in an overlapping extension PCR reaction, first 5 cycles without primers (T_anneal_=30°C), then 30 cycles with primers (T_anneal_=50°C). This PCR product was used to introduce the D262N mutation by overlapping extension PCR. These WT and MUT PCR products, and a PCR-amplified *gfp* gene, were cloned into AvaI linearised pT2A_Ddi1_cKO using In-Fusion to generate pT2A_PfDdi1_cWT and pT2A_PfDdi1_cMUT.

For the conditional tagging of PfDdi1 with a single HA tag, the RR2 region was amplified from the pT2A_PfDdi1_cWT. A long reverse oligonucleotide containing the HA tag sequence was annealed to the end of the RR2 region for 5 cycles (T_anneal_=35°C). Primers were then added to the reactions to amplify RR2-HA, and the resulting amplicon was used in an In-Fusion reaction with AvaI digested pT2A_PfDdi1_cWT to give pT2A_PfDdi1_cHA.

### Culture and transfection of *P. falciparum* cultures

All *P. falciparum* cell lines were cultured *in vitro* in human erythrocytes (UK Blood and Transfusion Service) at 37°C in RPMI-1640 (Gibco) based media supplemented with 5% (w/v) Albumax II and 2 mM L-glutamine. Synchronization was performed as previously described (Blackman, 1994). Cultures were monitored by light microscopy of methanol fixed and Giemsa (VWR International) stained thin-blood smears. Mature schizonts were collected by centrifugation on a Percoll cushion (GE Healthcare Life Sciences) (Kramer et al., 1982).

Transfection of *P. falciparum* 1G5DiCre (C. R. Collins, Das, et al., 2013) or B11 (Perrin et al., 2018) mature schizonts was performed as previously described (Das et al., 2015) using 10 μg plasmid DNA. 24h following transfection, media was replaced and supplemented with 2.5 nM WR99210 (Jacobus Pharmaceuticals) to select for a human dihydrofolate reductase cassette on the episome backbone. Once WR99210 resistant cultures reached 1% parasitaemia, cultures were treated with 450 μg/ml Geneticin Selective Antibiotic G418 Sulfate (Gibco) (Birnbaum et al., 2017). After selection, cultures were maintained on 2.5 nM WR99210 and 225 μg/ml G418, and parasites cloned by plaque assay (Thomas et al., 2016). Parasite cultures were serially-diluted across a flat-bottomed 96-well plate (Corning Costar) at a haematocrit of 0.75% and incubated for 12 days before single plaque-containing wells were selected as clones.

Unless otherwise stated, synchronous ring stage *P. falciparum* cultures were treated for 3h with 100 nM RAP (Sigma-Aldrich) or DMSO at 37°C (C. R. Collins, Das, et al., 2013). The cultures were then washed with RPMI media before the addition of drug-free complete media. To evaluate integration and excision by PCR, gDNA was extracted from saponin-lysed parasites using DNeasy Blood and Tissue Kit (Qiagen).

### Analysis of parasite samples by flow cytometry

Parasite samples to be analyzed by flow cytometry were fixed in 4% paraformaldehyde (Sigma) and 0.02% glutaraldehyde (Sigma) for 1h at room temperature (RT), diluted 5-fold in PBS, and stored at 4°C until analysis. Samples were incubated with 1 μg/ml of the DNA stain Hoechst-33342 (ThermoFisher Scientific) for 30 min at 37°C and analysed on a Fortessa flow cytometer (BDBiosciences). All erythrocytes were gated depending on their forward and side scatter profiles and iRBCs identified as the DNA-positive population using a UV laser (355 nm) and a 450/50nm emission filter. To study RNA content of iRBCs, samples were also stained with 2 μM of the 132A RNA dye (kind gift of Prof Young-Tae Chang, University of Singapore) (Cervantes et al., 2009), and the signal detected using a blue laser (488 nm) and a 610/20 nm emission filter. To measure GFP signal, the blue laser (488 nm) was used with a 525/50nm emission filter. Analysis of flow cytometry data was carried out in FlowJo software (version 10).

### Plaque assay

DMSO or RAP treated trophozoite stage parasites were diluted to approximately 250 parasites/ml in 0.75% haematocrit, and 30 wells (200 μl) were plated per condition in 96 flat-bottomed well plates. Plates were incubated at 37°C for 12 days prior to scanning the plates with a Perfection V750 Pro scanner (Epson). Plaques were counted using an automatic counting protocol. Wells were selected in PhotoShop (Adobe) with the Magic Wand tool (tolerance of 75) to remove the background of the 96-well plate from further analysis. The TIFF images of the selected wells were opened in Fiji (Schindelin et al., 2012). The RGB image channels were split in order to obtain the green channel image, which was was binarized (black on white) so that plaque number and size could be counted using the Fiji ‘Analyse Particles’ function, with a size cut-off of 20-750 pixels and a circularity of 0.5-1.00.

### Replication and invasion assays

DMSO or RAP treated trophozoite-stage cultures were diluted to 0.1% parasitaemia in 0.5% haematocrit, plated in triplicate (200 μl) in a round-bottomed 96 well plate, and cultured for three cycles with media replaced at 120 hpi. Samples were fixed for FACS analysis at 24, 72, 120 and 186h following DMSO or RAP treatment. For invasion assays, schizonts were Percoll purified and added to fresh erythrocyte suspensions at 1% haematocrit and 3% parasitaemia. Samples were plated in triplicate (200 μl) into round-bottomed 96 well plates and incubated at 37°C overnight. Samples were fixed at 0 and 24 h and invasion rates determined by flow cytometry. To determine the effect of C2 (MRC Technology) (C. R. Collins, Hackett, et al., 2013) treatment on invasion efficiency, DMSO or RAP treated parasites were treated with 1 μM C2 or DMSO at 40 hpi, washed with media at 52 hpi, and cultured for an additional 4h before fixation. Samples were analyzed by flow cytometry to determine the number of rings and schizonts present in the cultures.

### Treatment of *P. falciparum* cultures with proteasome inhibitors

DMSO or RAP treated parasite lines at 2% parasitaemia and 1.5% haematocrit were treated with different concentrations of BTZ or CPZ (Sigma) at 36 hpi. BTZ was washed out from the samples after 4h of treatment, and parasites cultured for an additional 24h. CPZ treated parasite were cultured without washing away the drug. At 58 hpi, parasites were fixed, and parasitaemia quantified by flow cytometry.

### Preparation of protein lysates for SDS-PAGE

Parasite samples were lysed for 5 min in 0.15% saponin to lyse the RBC and PV membranes at RT followed by centrifugation at 17000 x g for 5 min. After washing in PBS, the parasite pellet was snap frozen and stored at -80°C. Schizont samples were Percoll-purified prior to saponin-lysis. Frozen pellets were thawed into 10 volumes of ice-cold hypotonic lysis buffer (0.1% SDS, 1% Triton X-100, 1x cOmplete protease inhibitor cocktail (Roche)), incubated on ice for 20 min, and centrifuged (17000 x g, 20 min, 4°C). The supernatant was combined with 4x Laemmli buffer containing 400 mM DTT, boiled for 5 min, and 10 μg of protein subjected to SDS-PAGE.

### Western blot analysis

A list of primary and secondary antibodies used in this study for WB and IFA are listed in Table supplement 2. Protein gels were transferred onto nitrocellulose membranes and subsequently stained with Ponceau S (Biotium) before blocking in 5% BSA (Sigma) in PBS with 0.1% Tween-20 (PBST). Blots were incubated with primary antibodies and subsequently horseradish-linked secondary antibodies, which were diluted in 2% BSA in PBST. For anti-HA probing, a biotinylated secondary antibody was used and was followed by a third incubation step with a streptavidin-linked horseradish peroxidase (Sigma), followed by washing 3 x 5 min with PBST. Visualization of HRP signal was achieved using Immobilon Western Chemiluminescent HRP Substrate (Millipore) and a ChemiDoc Imager (Bio-Rad).

### Fluorescence microscopy

Thin smears of infected erythrocytes were air dried and were fixed in 4% formaldehyde for 30 min and permeabilized in 0.1% Triton X-100 (Sigma) for 10 min. The slides were blocked in 3% BSA in PBS for at least 1 h. All antibodies used in IFA were diluted with 3% BSA in PBS. Slides were probed with primary antibodies followed by AlexaFluor antibody (Invitrogen). In the case of anti-HA probing, a 3-step approach was taken using a biotinylated secondary antibody and a final incubation step with a strepatividin-linked AlexaFluor antibody (Invitrogen). Slides were inserted into a 1 μg/ml solution of 4’,6-diamidino-2-phenylindole (DAPI, Sigma) for 10s and, after washing, were mounted with Citifluor anti-fade mounting medium. Sealed slides were analyzed on a Leica TCS SP5 Confocal microscope using 100x oil immersion objective or a Nikon Eclipse Ni-E widefield upright microscope with a Hamamatsu C11440 digital camera and Nikon 100x oil immersion objective. Images were processed using Leica Application Suite Advanced Fluorescence, NIS Elements software (Nikon), and Fiji.

Fluorescence microscopy analysis of samples stain with the RNA dye 132A was performed as previously described (Bell, et. al, 2020). Briefly, parasite cultures samples were fixed in 4% paraformaldehyde (Sigma) and 0.02% glutaraldehyde (Sigma) for 1h at RT, diluted 5-fold in PBS and stored at 4°C. These were then stained with 2 μM 132A, 1 μg/mL Hoechst, and 0.2 μg/mL WGA-647, transfer to a flat-bottom glass-bottom 96-well plate and incubated for 30 min at 37°C, which is sufficient time to for the cells to at the bottom of the well. WGA-647 binds to lectin on the surface of the RBC. Images were collected on an Olympus IX83 inverted microscope using a PLAPON O 60/1.42 objective lens and a Hamamatsu ORCA-Flash 4.0 sCMOS camera (pixel size 6.5μm), and the Olympus CellSens Dimention acquisition software with well plate navigator (version 1.6). A drop of immersion oil was manually dispensed under each well and 25 images per well were collected in four channels: BF (differential interference contrast), Hoechst (λ_ex_ = 390/18 nm, λ_em_ = 440/40 nm), 132A (λ_ex_ = 560/25 nm, λ_em_ = 607/36 nm) and WGA-647 (λ_ex_ = 650/20 nm, λ_em_ = 692/40 nm). Images were analysed using Fiji.

### Serial block-face scanning electron microscopy (SBF-SEM)

PfDdi1_cMUT parasites were DMSO or RAP treated as standard, and schizonts purified at 44 hpi and treated with 1 μM C2 for 4 h. Schizonts were then washed in RPMI, and fixed with 10 volumes of 4% formaldehyde in 0.1 M phosphate buffer pH 7.4 (PB) for 15 min at RT followed by centrifugation (2400 x g, 1 min). A second fixative step was performed on the pelleted material with 2.5% glutaraldehyde and 4% formaldehyde in PB for 30 min at RT followed by centrifugation (2400 x g, 1 min). The pellet was resuspended and stored in 1% formaldehyde in PB at 4°C. Infected erythrocytes were embedded in 2% agarose, sliced into 1 mm^3^ sections, and washed in PB. Cells were incubated with 2% reduced osmium for 1 h at 4°C, washed in ddH_2_O (3 x 5 min for all ddH_2_O washes), incubated with 1% thiocarbohydrazide for 20 min at RT, washed in ddH_2_O, incubated in 2% aqueous osmium tetroxide for 30 min at RT, washed in ddH_2_O and incubated in 1% uranyl acetate at 4°C overnight. Cells were washed in ddH_2_O, incubated in lead aspartate for 30 min at 60°C and washed in ddH_2_O. Samples were subjected to dehydration by a graded ethanol series (70%, 90%, 100%). Samples were infiltrated with 50:50 propylene oxide: Durcupan for 1h at RT, then with 100% Durcupan twice for 2h at RT. The samples were transferred to fresh 100% Durcupan and baked at 60°C for 48h. A Sigma SEM (Zeiss) with integrated Gatan 3View was used to analyze resin blocks of infected erythrocytes by SBF-SEM using 50 nm slices.

### Pull-down of PfDdi1-HA and LC-MS/MS

PfDdi1_cHA parasites were DMSO or RAP treated, and mature C2-treated schizonts were saponin-lysed, washed, snap frozen, and stored at -80°C. The frozen schizont pellets (∼500 μl) were lysed in six volumes of ice-cold 20 mM Tris-HCl pH 7.6, 150 mM NaCl, 1% Triton X-100, 1 mM EDTA and 1 x cOmplete protease inhibitor cocktail (Roche) with three rounds of freezing in liquid nitrogen and rapid thawing. After 20 min on ice, the lysate was centrifuged (17000 x g, 20 min, 4°C) and the resulting supernatant further clarified by passing through a SpinX filter column (Corning Costar, 0.22 μm) at 17000 x g for 1 min at 4°C. The flow-through was added to 90 μl Anti-HA Magnetic Beads (Pierce) pre-washed in wash buffer (50 mM Tris HCl pH 7.6, 150 mM NaCl, 0.05% Tween-20) and incubated overnight at 4°C on a rotating mixer (VWR). A Dynabeads MPC-S Magnetic Particle Concentrator was used to collect the magnetic beads and extract the unbound material. The beads were washed in wash buffer (5 x 300 μl) prior to boiling in 75 μl 1 x Laemmli buffer for 5 min. To monitor the purification, samples were taken throughout the process and were boiled in 1 x Laemmli buffer for 5 min. These samples and the eluted material were subjected to SDS-PAGE, followed by Quick Blue (triple red) and Silver Staining (Bio-Rad) or anti-HA WB to monitor pull-down.

For LC-MS/MS analysis, 50 μl of eluted samples were electrophoresed for 8 min at 200 V on a 10% Mini-PROTEAN TGX Precast Protein Gel (Bio-Rad) and protein was stained using Quick Blue Stain. Samples were sliced into 2×2 mm gel pieces, trypsinised, the peptides extracted, dried using an SPD SpeedVac (Thermo Scientific Savant) and resuspended in 0.1% trifluoroacetic acid (TFA). In triplicate, peptides were separated on a 50 cm, 75 μm inner diameter Easyspray column over a 120 min gradient and eluted onto an Orbitrap mass spectrometer (ThermoFisher Scientific). Xcalibur software (ThermoFisher Scientific) was used during data acquisition. Data analysis was performed in MaxQuant software, using *P. falciparum* PlasmoDB and *H. sapiens* Uniprot databases. Label-free quantification results were transferred to Perseus (version 1.4.0.11) (Tyanova et al., 2016) for data transformation.

### Processing of hyper-ubiquitinated proteins

Parasite samples were treated with 0.15% saponin for 5 min, washed twice in PBS, frozen in liquid nitrogen, and stored at -80°C. Upon thawing, samples were incubated for 20 min in 50 mM Tris-HCl pH 7.4, 2 mM EDTA, 150 mM NaCl, 1% Triton X-100, 2 mM N-Ethylmaleimide, 2.2 mM PMSF, 2 mM benzamidine HCl, 2 μM Leupeptin and 1 μg/ml pepstatin A. The lysates were sonicated at 20% amplitude for 15 seconds (Branson digital sonifer model 250) and centrifuged 17000 x g for 5 min. The soluble material was retained, and 15 μg of each sample was fractionated by SDS-PAGE on Criterion TGX 4-15% gels (BioRad). WB was performed using P4D1 anti-ubiquitin antibody (Santa Cruz Biotechnology). Pre-treatments with 6 μg of recombinant *P. falciparum* Ddi1, hDdi2, or LmDdi1 protein were carried out at 30°C for 30 min prior to SDS-PAGE and WB analysis.

### Recombinant expression of PfDdi1

To express full-length PfDdi1, we used a recently published approach that uses a SUMO tag to aid solubilization of the protein of interest (Kuo et al., 2014) and that was successfully used to produce recombinant hDdi2 and *Leishmania* Ddi1 in *E. coli*. The SUMO tag can be cleaved using the Ulp1 SUMO protease. PfDdi1 was expressed in *E. coli* with an N-terminal His_6_, SUMO and FLAG tags, and purified using a 4-step approach including a step to remove the N-terminal SUMO tag. A catalytically inactive D262N mutant was also expressed in this way.

The entire Ddi1 expression construct was ordered from GenScript and contained the *PfDdi1* gene recodonised to *E. coli* alongside sequences encoding an N-terminal hexahistidine, SUMO and FLAG tags. The PfDdi1 D262N mutant was generated by site-directed mutagenesis with overlapping extension PCRs. These inserts were cloned into the NdeI and BamHI digested pET11a vector giving pET11a_6His-SUMO-FLAGPfDdi1 or pET11a_6His-SUMO-FLAG-PfDdi1-D262N.

Expression constructs were transformed into BL21 (DE3) (New England Biolabs) bacteria in the presence of 100 μg/ml ampicillin. Single colonies were used to inoculate lysogeny broth with 100 μg/ ml ampicillin, and were incubated overnight at 37°C. This starter culture was used to inoculate 500 ml of lysogeny broth with 100 μg/ml ampicillin at 37°C. Once an optical density of 0.4-0.8 units at 600 nm was achieved, 0.1 mM IPTG (Sigma) was added, and cultures were incubated overnight at 16°C. Bacteria were pelleted and frozen at -20°C.

The frozen bacterial pellet was resuspended in 10 volumes of lysis buffer (1 x PBS pH 7.4, 20 mM imidazole, 0.5 mg/ml lysozyme, 1 x Protease Inhibitor Cocktail EDTA free Set V (CalbioChem)) for 30 min at RT. This lysate was sonicated for 2 min (15 s on and 20 s off for 2 min at 20% amplitude) and centrifuged at 100,000 x g for 40 min at 4°C (Avanti J-301 Beckman Coulter centrifuge with rotor JA-30.50). The supernatant was filtered and passed through preequilibrated 5 ml HisTrap FF columns (GE Healthcare), washed with 5 columns volumes (CVs) of PBS pH 7.4 supplemented with 450 mM NaCl and 70 mM imidazole, and eluted with 10 CVs of PBS pH 7.4 containing 150 mM NaCl and 500 mM imidazole. The fractions with the purest protein were pooled and concentrated to <10 ml using a 3 Molecular Weight Cut Off (MWCO) centrifugal filter device (Amicon). 1000 units of SUMO Protease 1 (EZCut) was added in a Pur-A-Lyze dialysis tube (Sigma, 3.5 MWCO), which was incubated at 4°C overnight in 1 L of SUMO cleavage buffer (50 mM Tris HCl pH 8, 100 mM NaCl, 10 mM DTT). Size exclusion chromatography was performed to separate FLAG-Ddi1 from the cleaved SUMO protein or the full-length His-SUMO-FLAG-Ddi1 using a HiLoad 26/600 Superdex 200 pg column (GE Healthcare) in a buffer of 20 mM Tris HCl pH 7.5, 100 mM NaCl. A final anion exchange purification step was performed using a Resource Q 6 ml column (GE Healthcare), using 50 mM Tris HCl pH 8, 150 mM NaCl as the loading buffer, and a linear elution gradient of 150 mM to 1M NaCl over a 25 CVs. Fractions with the purest protein were combined and diluted in protein storage buffer (20 mM Tris HCl pH 7.5, 100 mM NaCl, 5% glycerol). Following SUMO cleavage, the mutant FLAG-Ddi1-D262N protein was purified first with anion exchange, then by size exclusion, before dilution in protein storage buffer. Protein was concentrated using a centrifugal device (Amicon, 3MWCO), frozen in liquid nitrogen, and stored at -80°C. Native mass spectrometry and circular dichroism were used to assess the protein quality (Figure S5).

### Circular dichroism analysis of recombinant protein

Purified FLAG-PfDdi1 and FLAG-PfDdi1^D262N^ were diluted to 0.2 mg/mL in 20 mM Tris-HCl, 1mM NaCl, and 0.05% glycerol, pH 7.5. Far UV circular dichroism spectra recorded from (190-260 nm) using a ChiraScan spectrophotometer (Applied PhotoPhysics) at 30 nm/min and a 0.1 cm path length.

### Intact protein MS analysis of recombinant protein

Fifteen picomoles of purified FLAG-PFDdi1 or FLAG-PfDdi1^D262N^ in 200 mM ammonium acetate were injected onto a Zorbax Porpshell 300SB-C3 column, washed with 0.1% formic acid in 5% acetonitrile, and eluted with 0.1% formic acid in 90% acetonitrile at 0.5 ml/min. Protein samples were ionised by electrospray ionisation in a Micromass Q-ToF API-US mass spectrometer. Mass estimation was using the MaxEnt1 algorithm of MassLynx software (version 4.1).

## Acknowledgements

We would like to thank Prof M. Blackman and his group for their support, assistance and suggestions about this work, and for providing antibodies and the 1G5 and B11 parasite cell lines. In particular we would like to thank F. Hackett and Dr C. Collins for assistance in parasite culture and Dr K. Kousis for very insightful discussion. We would also like to thank Dr A. Holder for providing antibodies, Dr. M. Treeck for the pT2A_FIKK10-cKO plasmid, and Prof Y.T. Chang for the 132A RNA dye. We are grateful to all members of the GlaxoSmithKline-Crick LinkLabs and a number of GlaxoSmithKline scientists for their guidance including Dr A. Bridges, Dr E. Jones, Dr R. Sundaresan, Dr Z. Nagurnaja and Dr J. Hutchinson. We are also grateful to Dr S. Kunzelmann of the Crick Structural Science Technology Platform for advice on circular dichroism. This work was supported by a BBSRC and GlaxoSmithKline Industrial Case Partnership Studenship (BB/N503678/1), by a Welcome Trust and Royal Society Sir Henry Dale Fellowship 099950. This work was also supported by the Francis Crick Institute which receives its core funding from Cancer Research UK (FC001043 and FC001166), the UK Medical Research Council (FC001043 and FC001166), and the Wellcome Trust (FC001043 and FC001166), and by grants to J.Q.S. from the European Research Council (Agreement 693327), a Laureate grant from the Novo Nordisk Foundation (NNF19OC0055875), and a Chair grant from the Danish National Research Foundation (DNRF153). For the purpose of open access, the authors have applied a CC BY public copyright licence to any author accepted manuscript version arising from this submission.

## Figures and Tables Supplements

**Table supplement S1.**
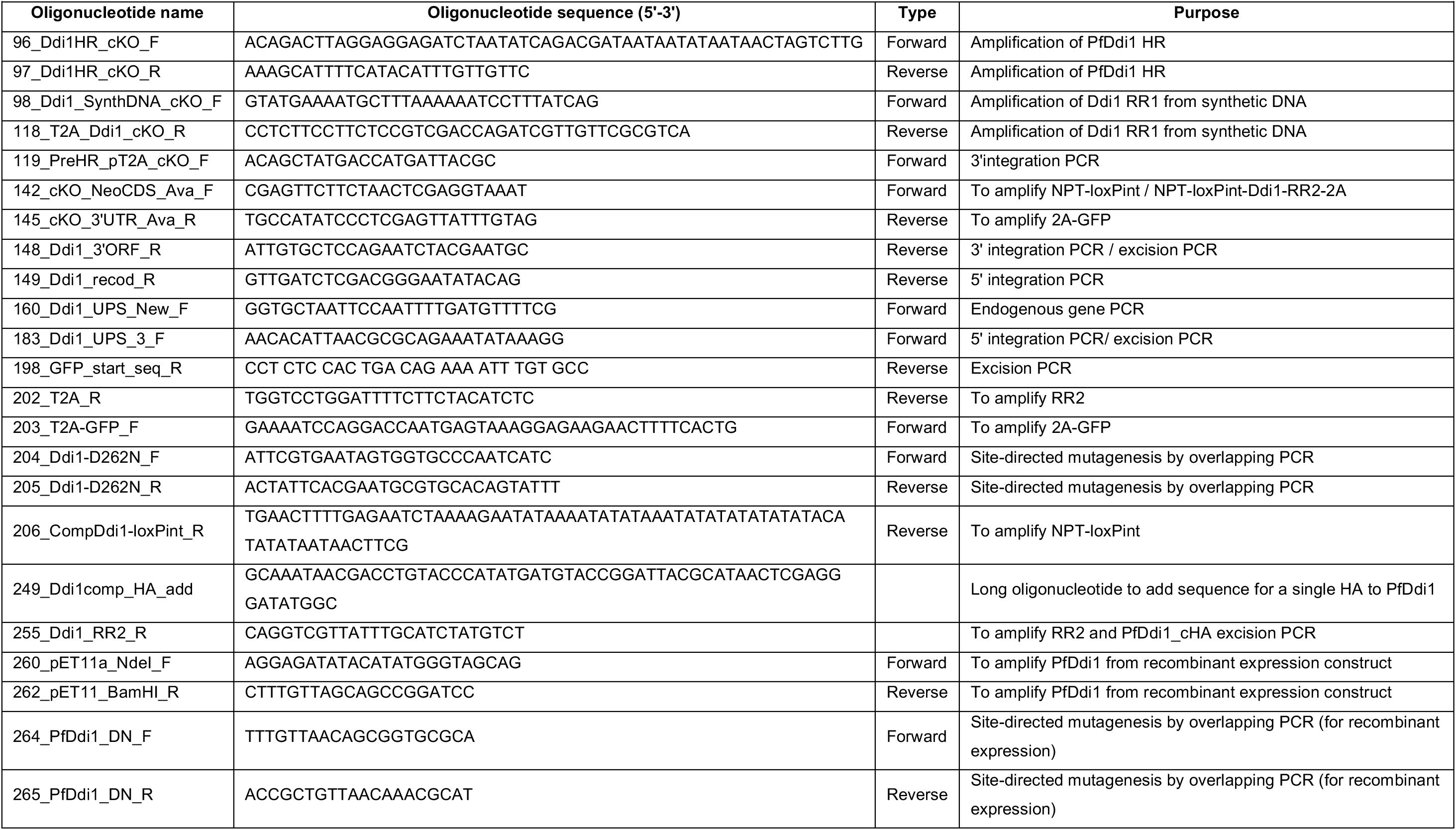
List of primers used in this study.

**Table supplement S2.** List of antibodies used in this study.

| **Table supplement S2. List of antibodies used in this study.** |  |  |  |  |  |
| --- | --- | --- | --- | --- | --- |
| **Antibody Name** | **Type** | **Animal** | **Use** | **Working concentration** | **Source** |
| Anti-HA 3F10 | Monoclonal | Rat | WB/ IFA | 1:1000, 1:100 | Roche |
| Anti-2A Peptide | Polyclonal | Rabbit | WB | 1:2000 | Merck Millipore |
| Anti-GFP (clones 7.1 and 13.1) | Monoclonal x2 | Mouse | WB | 1:1000 | Roche |
| Anti-His Tag (His.H8) | Monoclonal | Mouse | WB | 1:2000 | ThermoFisher Scientific |
| Anti-AMA1 | Polyclonal | Rabbit | WB/ IFA | 1:1000, 1:200 | Collins et al., 2009 |
| Anti-MSP1 (X-509) | Monoclonal | Human | IFA | 1:1000 | Blackman et al., 1991 |
| Anti-SERA5 | Polyclonal | Rabbit | WB/ IFA | 1:2000, 1:500 | Collins et al., 2017 |
| Anti-GAP45 | Polyclonal | Rabbit | IFA | 1:10000 | A. Holder, The Francis Crick Institute |
| Anti-EXP2 7.7 | Monoclonal | Mouse | IFA | 1:20 | Hall et al, 1983 |
| Anti-EBA-175 | Monoclonal | Rabbit | WB/IFA | 1:1000, 1:200 | Reed et al, 2000 |
| Anti-RhopH2 (mAb 61.3) | Monoclonal | Mouse | IFA | 1:5000 | Holder et al, 1985 |
| Goat anti-rabbit HRP | Polyclonal | Goat | WB | 1:10000 | Bio-Rad |
| Goat anti-mouse HRP | Polyclonal | Goat | WB | 1:10000 | Bio-Rad |
| Goat anti-rat biotin | Polyclonal | Goat | WB/ IFA | 1/8000. 1:1000 | ThermoFisher Scientific |
| Goat anti-rabbit Alexa 594/ 488 | Polyclonal | Goat | IFA | 1:2500 | Molecular Probes |
| Goat anti-human Alexa 594/ 488 | Polyclonal | Goat | IFA | 1:2500 | Molecular Probes |
| Goat anti-mouse Alex 594/ 488 | Polyclonal | Goat | IFA | 1:2500 | Molecular Probes |

**Figure 1-figure supplement 1.**
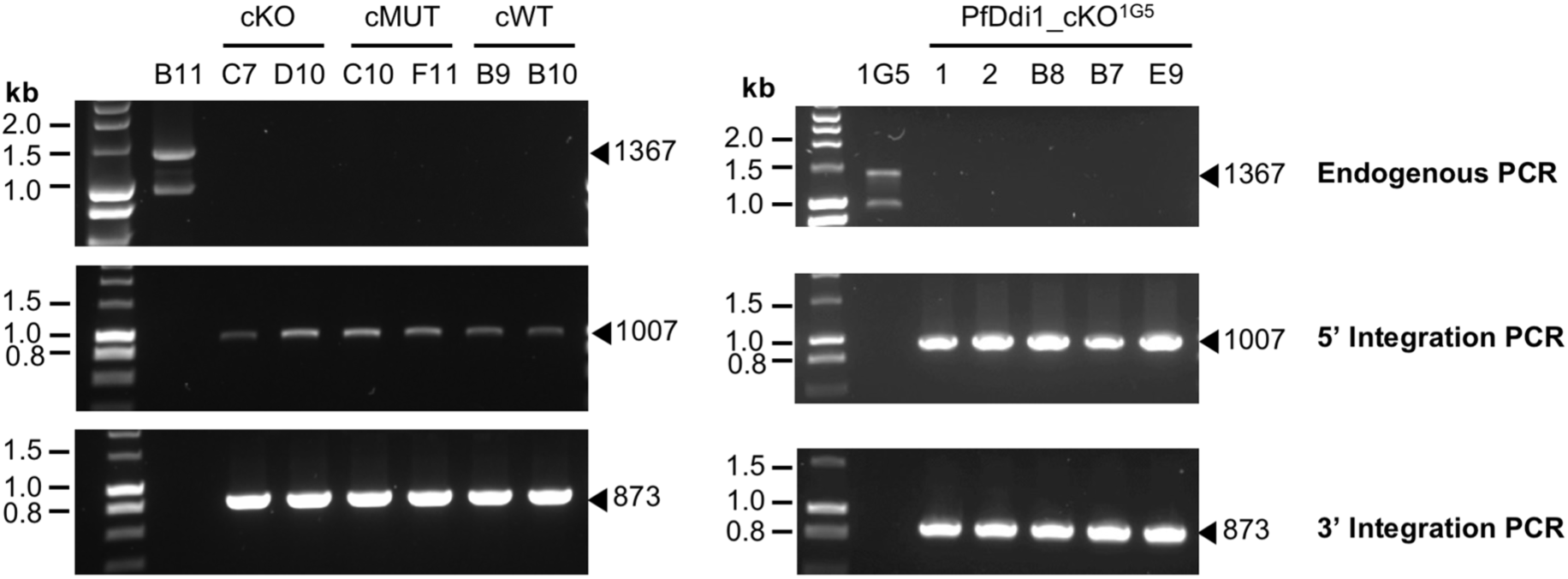
Integration PCRs. Integration of our cKO construct in the 1G5 line (right) and of the cKO, cMUT and cWT constructs in the B11 one (left) was confirmed by PCR. Diagnostic PCR for 3’-end and 5’-end integration was performed on genomic DNA from two clones of the PfDdi1_cKO, PfDdi1_cMUT, and PfDdi1_cWT lines, and for three clones of the PfDdi1_cKO^1G5^ line. Diagnostic PCRs were also performed before cloning of the PfDdi1_cKO^1G5^ lines (lanes 1 and 2 refer to two different transfections). Diagnostic PCR was also performed to check for the loss of the endogenous locus after integration. The size of the expected PCR products is shown with arrowheads. Genomic DNA from the B11 and 1G5 were used as negative controls.

**Figure 1-figure supplement 2.**
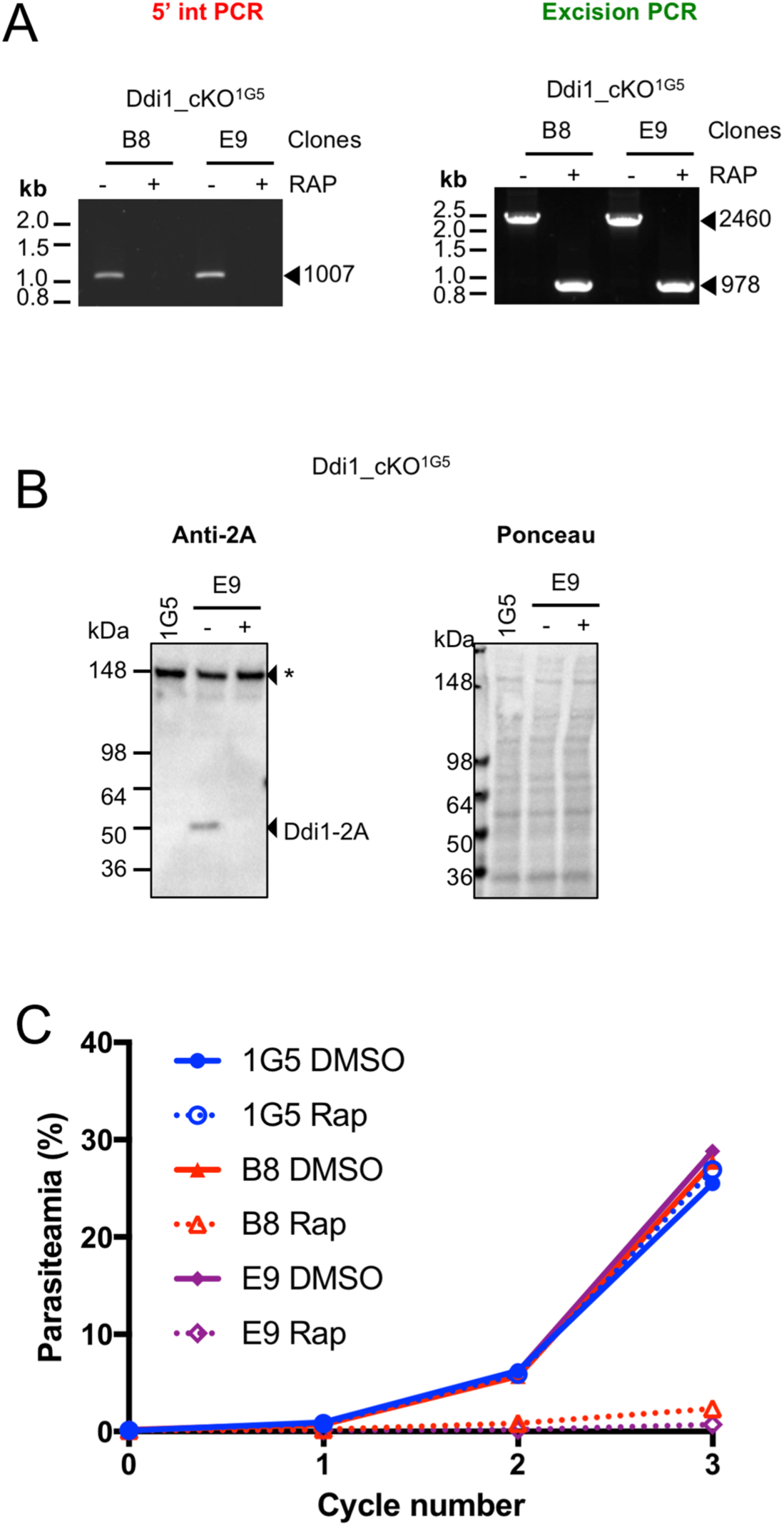
Characterization of the PfDdi1_cKO line generated in the 1G5 background line. (**A**) Diagnosis PCRs showing excision at the *PfDdi1* locus of two PfDdi1_cKO clonal lines. Genomic DNA was collected from the indicated lines 24h after DMSO or RAP treatment and used to perform diagnostic PCRs. The sizes of the expected PCR products are shown with arrowheads. RAP treatment of our conditional lines results in the loss of the 3’ primer binding sites in our integration PCR and a significant decrease in size of the excision PCR product. (**B**) WB analysis of the PfDdi1_cKO line. DMSO and RAP treated parasite lysates were collected at schizont stage and analyzed by WB using an anti-2A antibody. Lysates from the 1G5 line were used as a negative control. Ponceau staining of the blot is shown to control for protein loading. (**C**) Replication assay. Two PfDdi1_cKO were DMSO or RAP treated at ring stage, diluted to 0.1% parasitaemia, and the parasitaemia quantified over the following three cycles by flow cytometry. RAP treatment in a complete block of parasite replication.

**Figure 3-figure supplement 1.**
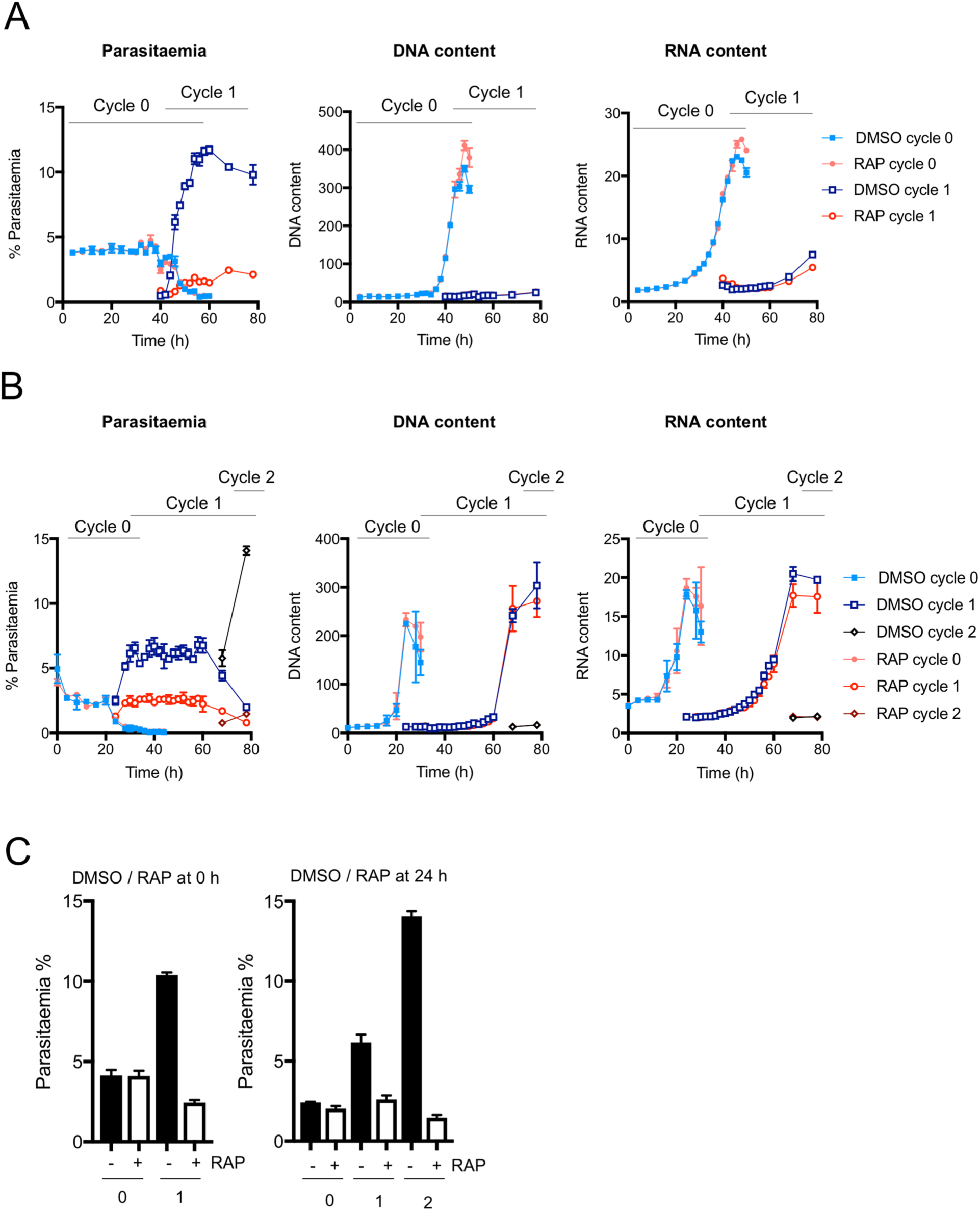
Effect of PfDdi1 KO on the erythrocytic cycle. (**A-B**) A tightly synchronized culture of PfDdi1_cKO was treated with DMSO or RAP at 0 hpi (**A)** or 24 hpi (**B**) and cultured for 80h. An aliquot of each culture was collected at different time points, fixed, and stored for FACS analysis. All samples were stained with Hoechst and the 132A dye to quantify the level of DNA and RNA signal in iRBC, respectively. At each time point, DNA content was used to differentiate iRBCs from uninfected RBCs (uRBCs). At the time of egress and invasion, between cycles 0 and 1 (or cycles 1 and 2), the level of DNA and RNA content was used to differentiate schizonts from rings, and thus assign which iRBCs belong to each cycle. For each population of iRBCs, parasitaemia was quantified (left), as well as the DNA (middle) and RNA (right) levels relative to the uRBCs population. DMSO and RAP treated samples are shown in blue and red, respectively. Independently of the time of RAP treatment, no changes in DNA or RNA content was evident in either cycle, suggesting that intracellular development is not affected by the loss of PfDdi1. Also, the gradual decrease in parasitaemia at the end of the cycle of treatment, indicative of schizonts egressing, is not significantly different between DMSO and RAP treated parasites. However, RAP treatment results in a very significant in the number of newly iRBCs after egress, strongly suggesting an invasion defect. **C)** Effect in parasite replication depending on the time of RAP treatment. For each cycle and treatment, parasitaemia was averaged over the time points where we did not observe egress or invasion. RAP treatment at 0 hpi results in a 6-fold decrease in parasitaemia in cycle 1 compared to DMSO-treated parasites, but only a 2-fold decrease when RAP treatment was performed at 24 hpi. This difference is presumably due to more PfDdi1 being present prior to gene disruption at 24 hpi compared to 0 hpi. While RAP treatment at 24 hpi resulted in no significant change in parasitaemia between cycles 0 and 1, we observed a 50% decrease in parasitaemia between cycles 1 and 2. This is consistent with what we observed between cycles 0 and 1 when parasites were treated at 0 hpi.

**Figure 3-figure supplement 2.**
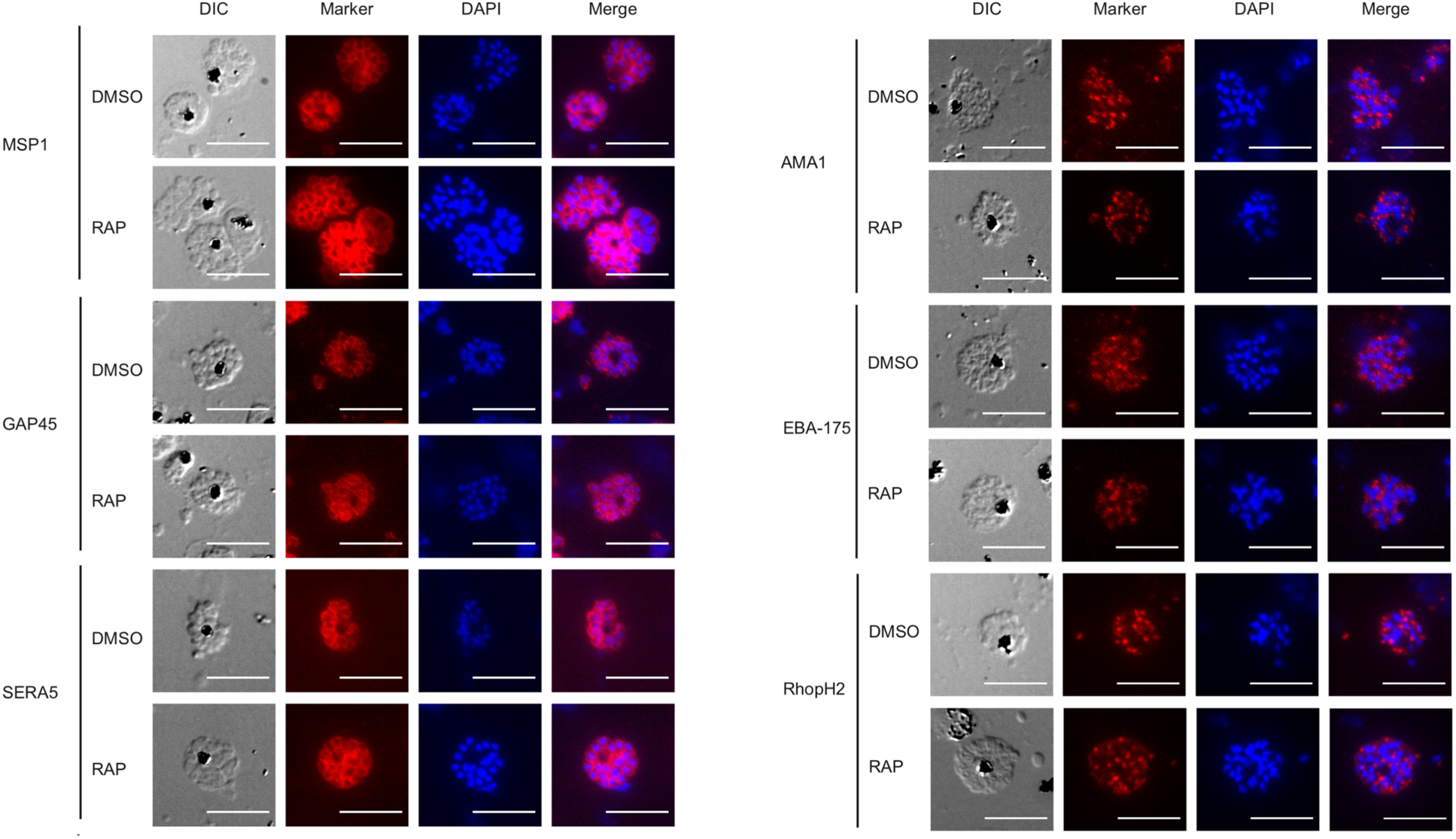
IFA studies. IFA of rhoptry, microneme, merozoite surface and PV proteinsshow the same localisation between wildtype and Ddi1-D262N schizonts. IFA analysis was performed on thin blood smears of DMSO or RAP treated PfDdi1_cMUT parasites that had been tightly synchronised and arrested immediately before egress by treatment with C2, i.e. mature segmented schizonts. Slides were probed with antibodies against MSP1 (merozoite surface), GAP45 (IMC), SERA5 (PV), RhopH2 (rhoptries), and AMA1 and EBA-175 (micronemes). Slides were also stained with DAPI to visualise parasite nuclei. Representative images are shown. Scale bar, 10 μm.

**Figure 4-figure supplement 1-7.** Videos showing each section image obtained by SBF-SEM. For Figure 4-figure supplement 1-3, the field is 20.48 x 20.48 microns. For Figure 4-figure supplement 4-7, the field is 32.77 x 32.77 microns.

**Figure 5-figure supplement 1.**
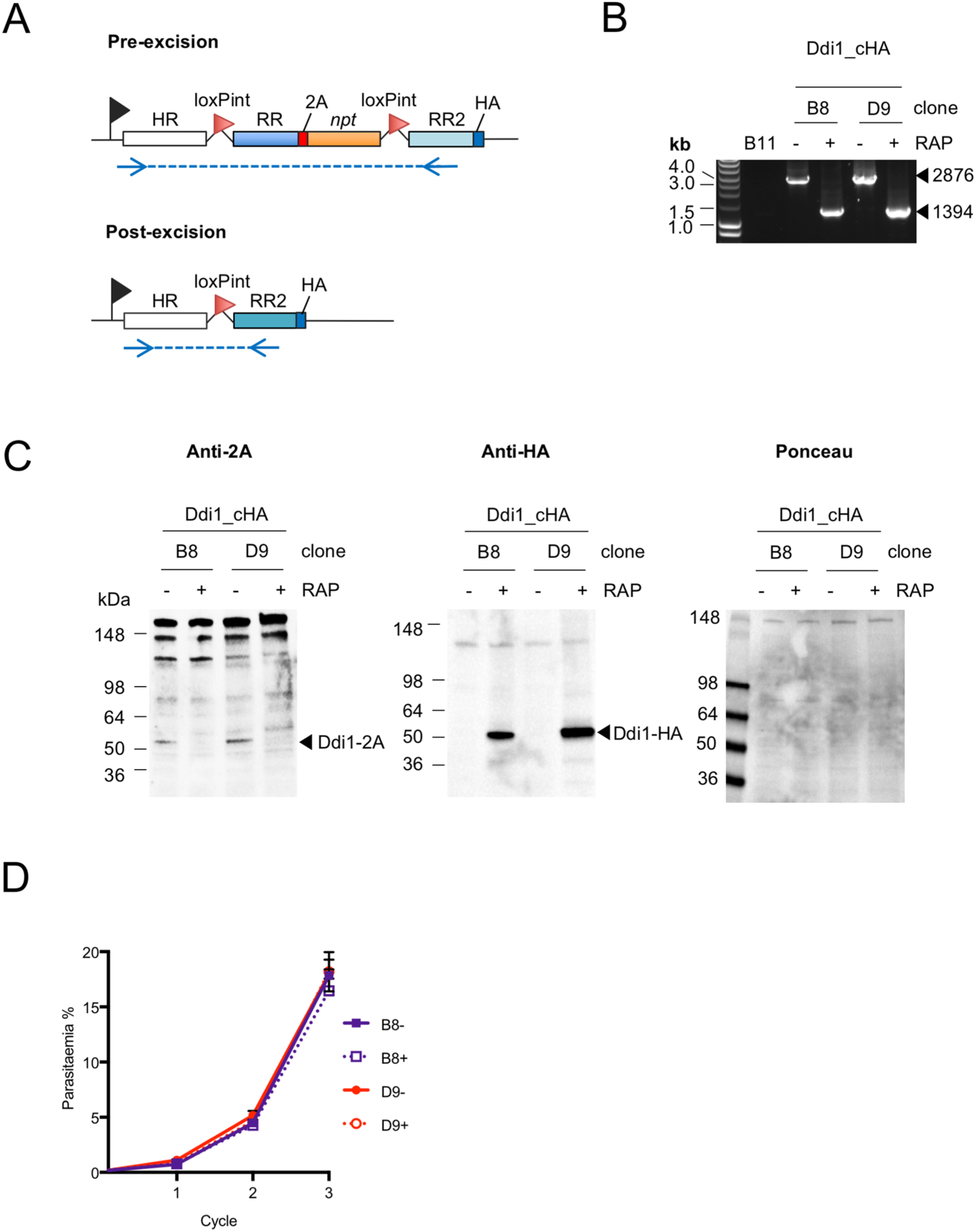
Characterization of the PfDdi1_cHA lines. (**A**) Schematic of the modified *PfDdi1* locus before (top) and after (bottom) RAP treatment. RAP treatment leads to excision of 2A-linked RR and *npt*, and replacement of the RR by RR2 that is C-terminally tagged with a single HA. Blue arrows indicate primer binding sites for excision diagnosis PCR. (**B**) Excision PCRs. RAP treatment of two PfDd1_cHA clones results in a decrease in the size of the excision PCR product. PCRs were performed after purification of genomic DNA 24h after DMSO or RAP treatment. The sizes of the expected PCR products are shown with arrowheads. (**C**) WB analysis of PfDdi1_cHA lines. The indicated clones were treated with DMSO or RAP at ring stage, and parasite lysates collected at schizont stage and analyzed by WB using either an anti-2A peptide (left) or anti-HA (middle) antibody. RAP treatment results in the loss of PfDdi1-2A and its replacement with PfDdi1-HA. Ponceau staining (right) is shown as a loading control. (**D**) Replication assay. Two PfDdi1_cHA clones were DMSO or RAP treated at ring stage, diluted to 0.1% parasitaemia, and the parasitaemia quantified over the following three cycles by flow cytometry. No differences in parasite replication was observed between DMSO or RAP treated cultures.

**Figure 6-figure supplement 1.**
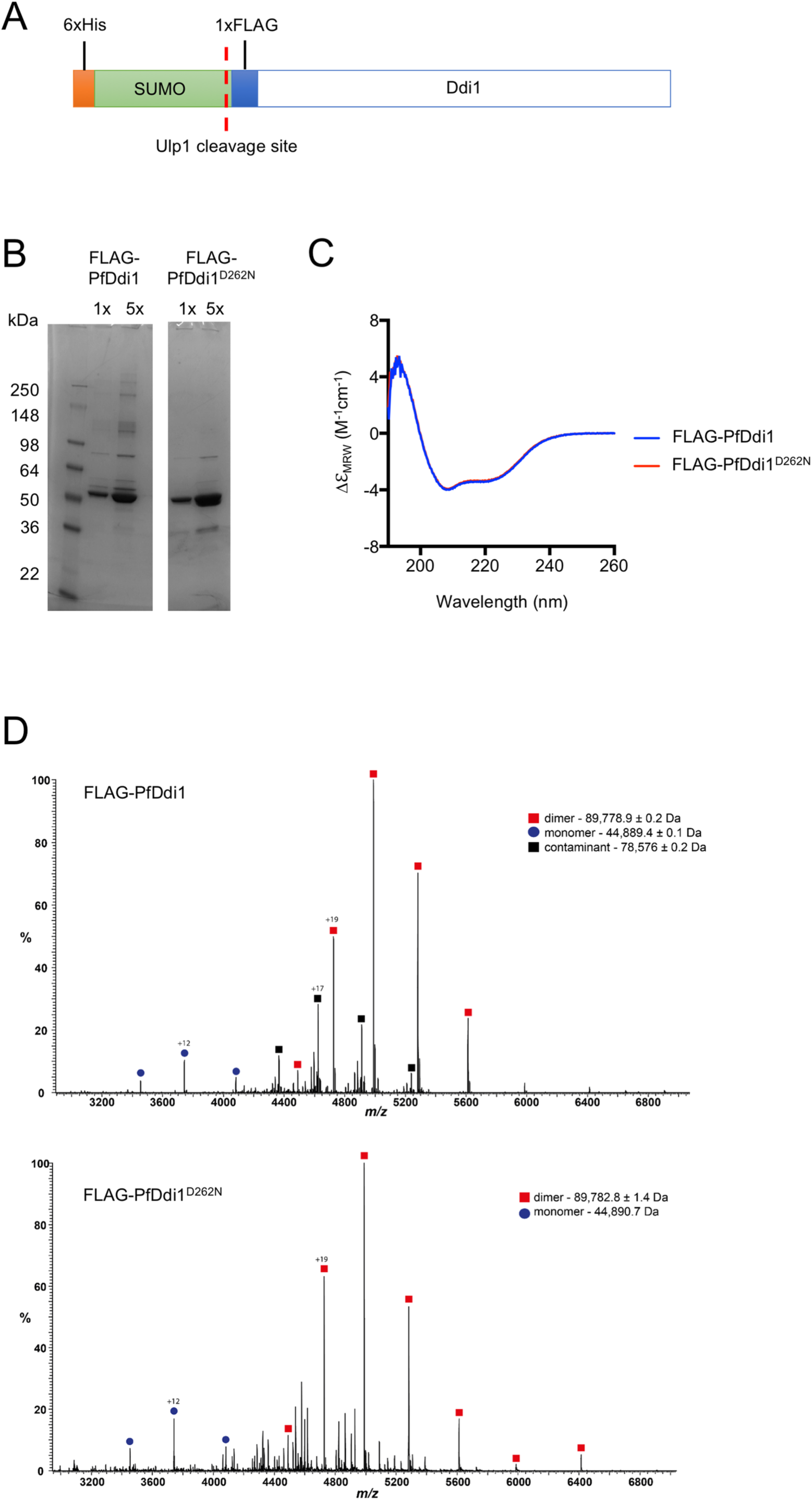
Quality control of recombinant PfDdi1. (**A**) Schematic of PfDdi1 expression construct. WT or MUT PfDdi1 was expressed with a 6xHis, SUMO and FLAG tag at the N-terminal. The SUMO tag improves solubilization and can be cleaved with the SUMO protease Ulp1. (**B**) Purity of recombinant protein. FLAG-PfDdi1 and FLAG-PfDdi1^D262N^ were purified using a 4-step purification method. Coomassie stained SDS-PAGE shows the purity of the obtained protein. For each protein 0.75 μM (1x) and 3.75 μg was loaded. (**C**). Circular dichroism analysis. The far UV spectra of FLAG-PfDdi1 and FLAG-PfDdi1^D262N^ obtained at 0.2 mg/mL. Both spectra are identical indicating that the D262N mutation does not disrupt the structure of PfDdi1. (**D**) Intact protein MS analysis of purified proteins. The predicted sizes of monomeric FLAG-PfDdi1 and FLAG-PfDdi1^D262N^ are 448911.16 and 44890.17 kDa, respectively, which is consistent with native mass spectrometry results. The distribution of monomeric (blue circles) versus dimeric (red circles) indicate that purified PfDdi1 is mainly homodimeric. Note that a contaminant (black squares) was detected in the FLAG-PfDdi1 sample. *m/z*, mass to charge ratio.

## Source Data Files Legends

**Figure 2-source data 1.** Related to Figure 2B. Table reporting the number of plaques counted in each plaque assay.

**Figure 2-source data 2.** Related to Figure 2C. Live microscopy of PfDdi1 KO parasites over 3 cycles. Synchronous PfDdi1_cKO parasites were RAP treated and highly mature schizonts were collected at the end of the cycle of treatment (Cycle 0) and of the following two cycles (Cycles 1 and 2). These samples were stained with DAPI and analyzed by live microscopy. Merged channels (DIC, GFP, and DAPI) of the microscope field used for quantification are shown.

**Figure 4-source data 1.** Related to Figure 4B. Merged fluorescence microcopy field images used for quantification of segmented vs. non-segmented schizonts after DMSO or RAP treated PfDdi1_cKO and 1G5 control parasites at 42, 44, and 46 hpi. RNA staining with the 132A dye is shown in red, DNA staining with Hoechst in blue, and RBC membrane staining with Alexa647-conjugated WGA in cyan. Only iRBCs containing schizonts (parasites with multiple nuclei) and hemozoin were used for quantification.

**Figure 5-source data 1.** Related to Figure 5D. Raw MS data of PfDdi1_cHA pull-down results. The excel file contains one sheet containing all identified proteins, and a second sheet showing only significantly enriched proteins in RAP-treated parasites relative to DMSO-treated ones.

